# Aetokthonotoxin, the causative agent of vacuolar myelinopathy, uncouples oxidative phosphorylation due to protonophore activity

**DOI:** 10.1101/2025.03.04.637646

**Authors:** Valerie I. C. Rebhahn, Mohamad Saoud, Mathias Winterhalter, Franziska Schanbacher, Maximilian Jobst, Rebeca Ruiz, Alexander Sonntag, Johannes Kollatz, Rieke Sprengel, Stephen F. Donovan, Giorgia Del Favero, Robert Rennert, Timo H. J. Niedermeyer

**Author notes:** **Corresponding author:** Prof. Dr. Timo H. J. Niedermeyer, **Email:**. **Author contributions:** V.I.C.R., M.S., M.W., F.S., M.J., R. Ruiz, A.S., J.K., G.D.F., R. Rennert, T.H.J.N. designed research, V.I.C.R., M.S., M.W., F.S., M.J., R. Ruiz, A.S., J.K., and R.S. performed research; V.I.C.R., M.S., M.W., F.S., M.J., R. Ruiz, G.D.F., R. Rennert and T.H.J.N. analyzed data; S.F.D. provided crucial insights, V.I.C.R. and T.H.J.N. wrote the paper with contributions from all other authors. **Competing Interest Statement:** The authors do not declare any competing interests.

## Abstract

Aetokthonotoxin (AETX) is an emerging environmental toxin produced by the freshwater cyanobacterium *Aetokthonos hydrillicola*. Accumulating in the food chain, it causes vacuolar myelinopathy, a neurological disease affecting a wide range of wildlife characterized by the development of large intra-myelinic vacuoles in the white matter of the brain. So far, the mode of action of AETX is unknown. After discovering that AETX is cytostatic and arrests cancer cell lines in G_1_-phase, metabolomic profiling of AETX-treated cells as well as an assessment of the physico-chemical properties of the compound suggested that AETX is a weakly acidic uncoupler of mitochondrial respiration. We confirmed this hypothesis by *in vitro* assays on mammalian cells, finding that AETX has the expected effects on the mitochondrial network morphology, mitochondrial membrane potential, and oxygen consumption rates, resulting in affected ATP generation. We confirmed that AETX is capable of transporting protons across lipid bilayers. In summary, we demonstrate that AETX is a protonophore that uncouples oxidative phosphorylation in mitochondria, the primary event of AETX intoxication.

**Significance statement:** Aetokthonotoxin (AETX) is an emerging cyanotoxin. Produced by the cyanobacterium *Aetokthonos hydrillicola*, it is transferred through the food chain, affects the nervous system, and eventually causes mortality in animals of various taxa. Our finding that AETX is an unspecific uncoupler of mitochondrial respiration implies that it might also be harmful for human health upon ingestion and trophic accumulation. First steps towards a full risk assessment are needed. An important aspect in this regard is the elucidation of the toxin’s mode of action. We anticipate our findings to be a starting point for the development of an adverse outcome pathway addressing the formation of vacuolar myelinopathy, expanding the significance of our results to the future risk assessment of other environmental neurotoxins.

## Introduction

First diagnosed after a mass mortality event of bald eagles (*Haliaeetus leucocephalus*) at DeGray Lake, Arkansas in 1994 (1), vacuolar myelinopathy (VM) has spread throughout the southeastern United States. Today, it is known that VM affects wildlife of various taxa apart from birds. These comprise fish, amphibians, and reptiles (2). VM is an eventually lethal disease characterized by widespread vacuolization in the white matter of the brain and spinal cord of affected animals with myelinated axons (2, 3).

In 2021, the causative agent of VM was discovered: aetokthonotoxin (AETX, Fig. 1). AETX is a naturally occurring toxin synthesized by the freshwater cyanobacterium *Aetokthonos hydrillicola* (2). AETX is an unusual pentabrominated biindole alkaloid with a 1,2’-biindole linkage and an indole-3-carbonitrile group, outstanding features for a natural product (4). *A. hydrillicola* colonizes the leaves of the invasive neophytic water plant *Hydrilla verticillata*, and wildlife consuming the leaves of the infested plant ingest the AETX producing cyanobacteria as well (2). From there, AETX is passed through the food chain (2, 5, 6). Also animals without myelin, like mollusks, crustaceans, and nematodes have been found to be affected by AETX (2). As intoxicated animals are debilitated and more likely to be preyed on, AETX poses a severe threat to already endangered predators such as the Florida snail kite (*Rostrhamus sociabilis*) (6). The mode of action (MoA) of AETX, however, is still unknown. Several structurally different compounds have been linked to the formation of similar edemas in the white matter of the brain (7). Despite their structural diversity, most of these compounds seem to interfere with energy metabolism, i.e., oxidative phosphorylation (7).

**Figure 1.**
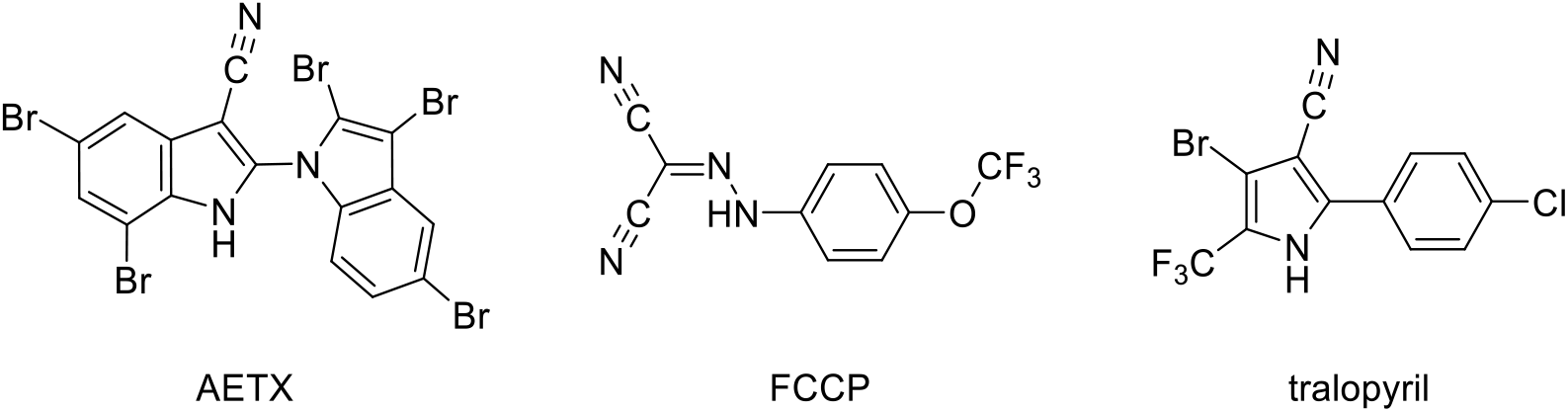
Structures of aetokthonotoxin (AETX), 2-[[4-(trifluoromethoxy)phenyl]hydrazinylidene]-propanedinitrile (FCCP), and tralopyril.

A multitude of methods have been developed in the past to elucidate the MoA of compounds, comprising, e.g., chemical proteomics techniques, affinity chromatography-based methods, phage display techniques, or yeast three-hybrid systems (8, 9). However, many of those methods have the drawback that they require derivatization of the compound of interest to enable, e.g., tagging with biotin or fluorophores (10). These modifications often affect the original interaction of the respective compound with its target (11). One technique that uses underivatized, genuine compounds is the recently described combination of metabolomics and machine learning for MoA prediction by pattern recognition. This mixed *in vitro* / *in silico* approach is based on the relative quantification of metabolites involved in central carbon and cellular energy metabolism, although downstream effects can also be detected in case compounds affect non-metabolic targets (12).

Using this method, we found that the metabolomic changes of AETX-treated cells clustered closely with those of cells treated with uncouplers of oxidative phosphorylation. In addition to a detailed characterization of the impact of AETX on cellular energy balance and *in vitro* oxygen consumption rate, we show that AETX can act as a protonophore on artificial lipid bilayer membranes. Structure-activity relationships (SAR) for AETX are discussed. Taken together, we confirm uncoupling of the oxidative phosphorylation as the MoA of AETX.

## Results & Discussion

### AETX is bacteriostatic in *Bacillus subtilis* and cytostatic in cancer cell lines, arresting them in G_1_-phase

Although the toxicity of AETX has been confirmed *in vivo* in birds (*Gallus gallus*), zebra fish (*Danio rerio*) and *Caenorhabditis elegans* (2), its activity on bacteria or mammalian cell lines has not yet been studied in detail. To assess its effect on bacteria, we evaluated the optical density of *Bacillus subtilis* and *Escherichia coli* suspensions after 20 h incubation with various AETX concentrations. While only a slight reduction of *E. coli* growth was observed at the highest concentration tested (30 μM), the minimal inhibitory concentration (MIC) of AETX for *B. subtilis* was 0.5 μM. However, even after 24 h incubation time with 30 μM, AETX had no bactericidal but bacteriostatic effect on *B. subtilis*. Darkfield microscopy of *B. subtilis* revealed a change of morphology from rod to spheric shape when treated with AETX at concentrations below the MIC (Fig. S1).

Toxicity of AETX for mammalian cells was tested by using a sulforhodamine B (SRB) cytotoxicity assay with the NCI-60 panel of the National Cancer Institute (NCI) Developmental Therapeutics Program (five-dose assay, 5 nM to 50 μM) (13), including colorectal cancer HCT116 and prostate cancer PC-3 cells, which were later also used by us for more detailed characterization. In addition, cytotoxicity was also determined in cervix carcinoma HeLa cells, and in fibroblast CCD1092Sk cells using the SRB assay. In most of the NCI-60 panel cell lines, AETX only showed moderate activity, with half-maximal lethal concentration (LC_50_) values of > 50 μM, while the half-maximal growth inhibition (GI_50_) was around 1 μM, and the total growth inhibition (TGI) around 5 μM (Fig. S2). We determined a half-maximal effect concentration (EC_50_) of 6.5 μM for HeLa cells (Fig. S3), which is in line with the aforementioned TGI determined by the NCI, and further agrees with our previously published data (14). Based on our results, the tested cancer cell lines seem to be more susceptible to AETX than the non-cancerous fibroblast cell line, as we were unable to determine an EC_50_ for the latter (Fig. S4).

Interestingly, visual examination of AETX-treated cells during routine brightfield microscopy of the assay plates revealed morphological changes of the cells treated with higher concentrations of AETX, rather than signs of cell death (Fig. S5). Along with the results of the SRB assays, the data indicated a cytostatic effect of AETX. To assess this hypothesis, we monitored the confluence of HCT116 cells before and after treatment with AETX. Indeed, we found a steady confluence level of cells treated with higher AETX concentrations starting at 5 μM AETX in contrast to control cells or cells treated with lower AETX concentrations (Fig. S6). This finding corresponds well to the TGI of HCT116 cells, as determined by the NCI.

To investigate this finding in more detail, we studied whether the cells were arrested in a certain phase of the cell cycle. Flow cytometric analysis of HCT116 cells treated for 24 h with solvent control, 1 or 5 μM AETX, and followed by DNA staining with propidium iodide, revealed an accumulation of the cells in G_0_/G_1_-phase for both AETX concentrations (Fig. S7). To further distinguish if cells accumulate in G_0_- or G_1_-phase, we stained HCT116 cells with a FITC-conjugated Ki-67 antibody. As only proliferating cells possess this marker (i.e., cells that are not in the G_0_-phase), a decrease in Ki-67 signal compared to the control would signify an arrest of cells in the G_0_-phase. As we found no significant difference between treated cells and the control (Fig. S8), we concluded that the cells were arrested in the G_1_-phase.

### Metabolomic profiling and physico-chemical properties suggest uncoupling as MoA of AETX

To generate a hypothesis about the potential MoA of AETX, its impact on the cells’ central carbon and energy metabolism was studied by metabolomic profiling of PC-3 cells incubated for 2, 4, 24, and 48 h with 0.74 μM AETX (EC_50_ in an MTT cell viability assay, Fig. S9). Using a previously published approach, the effects of AETX on the treated cells’ metabolome were compared with the effects of a panel of reference compounds with known MoAs (12).

When cells were treated for 2 h or 4 h, we found that the AETX treatment clustered hierarchically most closely with the outcome of treatments with [(3-chlorophenyl)hydrazono]malononitrile (CCCP), 2,4-dinitrophenol (DNP), and emodin, all classified as uncouplers of oxidative phosphorylation (12) (Fig. 2, Fig. S10). After 24 h and 48 h of treatment, AETX additionally clustered with hexachlorophene and bithionol (Fig. 2, Fig. S10), which, although they are classified as glutamate dehydrogenase (GDH) inhibitors in our model (12), are also uncouplers of oxidative phosphorylation (15, 16). In addition, adenosine monophosphate (AMP), the general marker of a reduced cellular energetic status, had a higher abundance in treated cells than in control cells (Fig. S11). In total, these findings underline the interference of AETX with cellular energy homeostasis and suggest uncoupling of the oxidative phosphorylation as the primary MoA of AETX.

**Figure 2.**
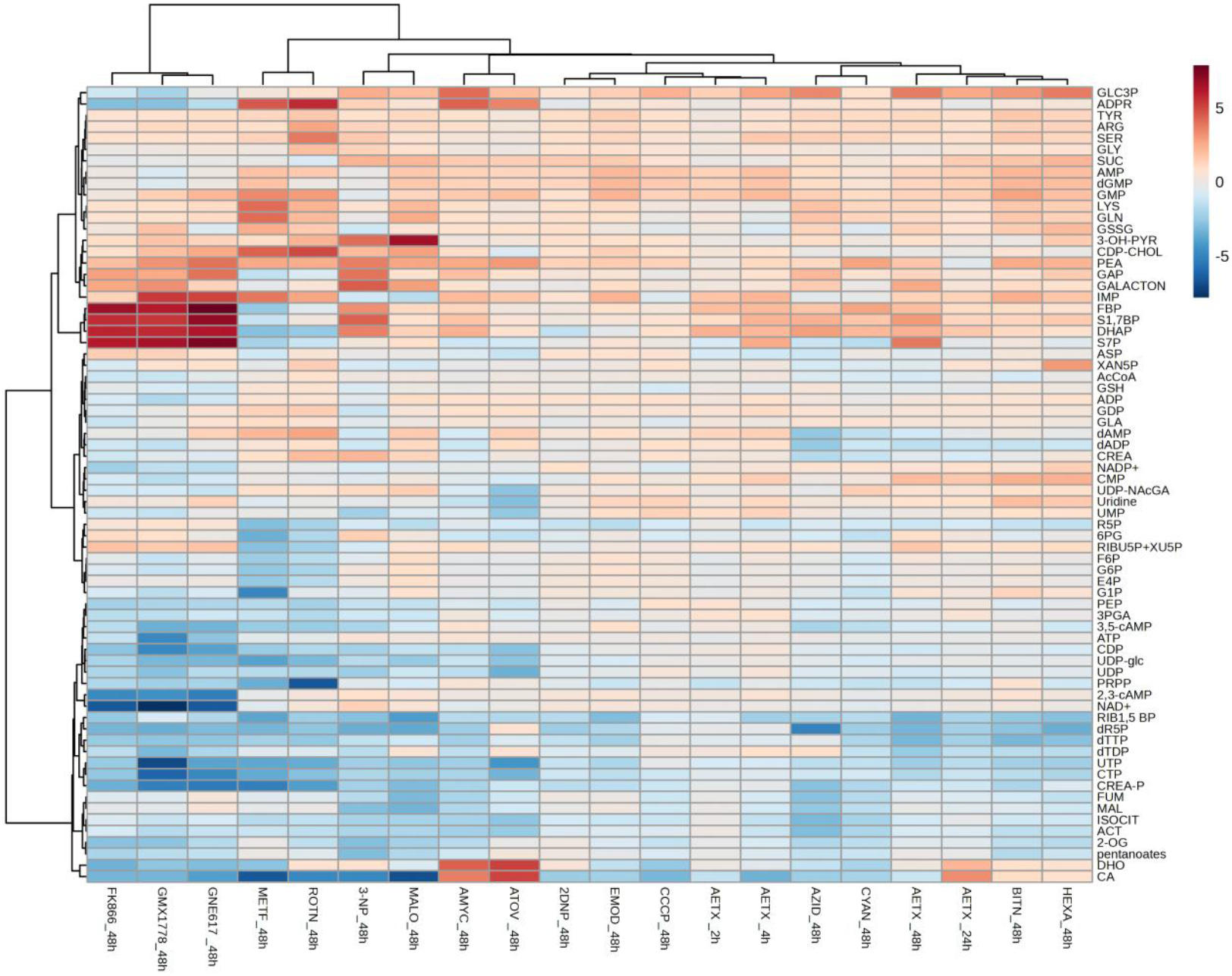
Relative abundance of key intermediates of the central carbon and cellular energy metabolism upon oxidative phosphorylation inhibition at complexes I-IV, treatment with uncouplers, NAMPT inhibition, GDH inhibitors and AETX treatment. The data represents the average log2-fold changes in peak areas which are relative to the cell number, after 48 h of reference drug treatment, and 2 h, 4 h, 24 h, and 48 h of AETX treatment (n=6), compared to a vehicle control (n=6). Compounds (mode of action): 2DNP - 2,4-dinitrophenol (uncoupler), 3-NP - 3-nitropropionic acid (CPLX II), AMYC - antimycin A (CPLX III), ATOV *-* atovaquone (CPLX III), AZID - sodium azide (CPLX IV), BITN *-* bithionol (GDH), CCCP - carbonyl cyanide chlorophenylhydrazone (uncoupler), CYAN - potassium cyanide (CPLX IV), EMOD *-* emodin (uncoupler), FK866 - FK866 (NAMPT), GMX - GMX1778 (NAMPT), GNE - GNE-617 (NAMPT), HEXA *-* hexachlorophene (GDH), MALO - malonic acid (CPLX II), METF - metformin (CPLX I), ROTN *-* rotenone (CPLX I); Abbreviations: CPLX I *-* complex I, CPLX II *-* complex II, CPLX III *-* complex III, CPLX IV *-* complex IV, GDH - glutamate dehydrogenase, NAMPT *-* nicotinamide phosphoribosyltransferase, Uncoupler *-* uncoupling of oxidative phosphorylation. For abbreviations of metabolites see supplementary metabolomics data (Excel file).

We subsequently examined the chemical structures and physico-chemical properties of known uncouplers. Many uncouplers, including CCCP and DNP, belong to a group called weakly acidic uncouplers. These molecules are in general characterized by high lipophilicity in conjunction with a weakly acidic functional group. Due to these properties, weakly acidic uncouplers can transport protons across biological membranes (protonophore activity), interfering, e.g., with oxidative phosphorylation in mitochondria (17, 18). Indeed, AETX perfectly matches these general requirements: Its pK_a_ value is 6.9 (Fig. S12), its logP is 4.7, and its logD at pH 7 is 4.4 (Fig. S13). According to Gange *et al*. (19), the optimum pK_a_ and logP values for uncoupling are 7.2 and 5.5, respectively, which is close to the values we determined for AETX. We further observed that the structure of tralopyril (Fig. 1), the active form of the prodrug chlorfenapyr, is strikingly similar to that of AETX. Both contain an aromatic NH group (indole in AETX, pyrrole in tralopyril) where the mesomeric effect of a nitrile group increases NH acidity, both feature an aromatic system adjacent to the NH-containing heterocycle, and both show halogen substitution that further increases NH-acidity due to electron withdrawal from the aromatic system and also increases lipophilicity. Approved as a biocide for restricted uses, chlorfenapyr exerts pan-species toxicity with a high vulnerability of birds (20). Trials in rodents showed similar effects and lesions in the brain and spinal cord as observed in animals with VM evoked from AETX (21). Likewise, the active metabolite of bromethalin, desmethylbromethalin, is an uncoupler that causes VM-like lesions in cat and bird brains (22, 23). Taken together, also our structural and physico-chemical evaluations support the hypothesis of uncoupling as MoA of AETX.

### AETX shows uncoupling-typical effects on mitochondria in mammalian cells

Mitochondrial bioenergetics is tightly linked to mitochondrial network morphology and *vice versa* (24). To validate the predictions discussed above *in vitro*, we studied how AETX affects the mitochondrial network in HCT116 cells. Immunofluorescence staining of the mitochondrial translocase of the outer membrane (TOM20) revealed a dose-dependent decrease of the TOM20 signal (Fig. 3A, 3B). Thus, AETX disrupts the mitochondrial network, visible already at the lowest concentration tested (0.1 μM). This result agrees with previous findings that compounds interfering with oxidative phosphorylation lead to mitochondrial fragmentation in most cell lines tested (25).

**Figure 3.**
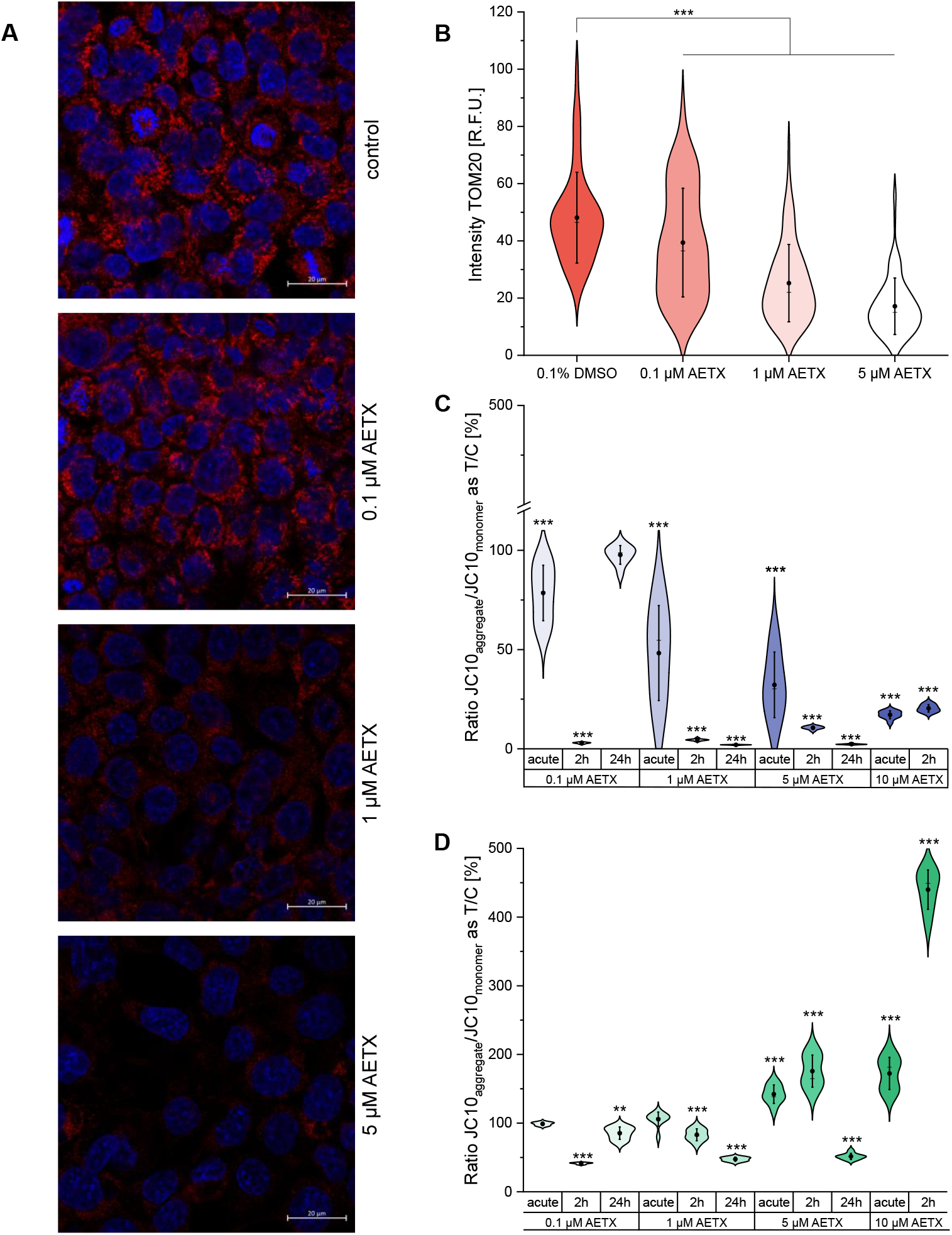
Impact of AETX on mitochondrial network and membrane potential. **(A)** Confocal microscopy images of immunofluorescence-stained translocase of the outer mitochondrial membrane 20 protein (TOM20) (red) and DNA (blue) in HCT116 cells. Cells incubated for 24 h with (from top to bottom) 0.1% DMSO (control), 0.1, 1, or 5 μM AETX. Scale bar 20 μm. Panels **B/C/D**: Violin plots with the mean indicated as black dot and the median as line. Standard deviation (1x) indicated as whiskers. Increasing shades of colour indicate increasing AETX concentrations. **(B)** Intensity of TOM20 signal as relative fluorescence units (R.F.U.) in HCT116 cells. Statistically significant difference to the control indicated with *** p < 0.001, obtained with Student’s t-test. Mitochondrial membrane potential measurement at different timepoints as ratio of JC10 aggregate to JC10 monomer in **(C)** HeLa cells or **(D)** fibroblasts normalized to control. Statistically significant difference to the control indicated with ** p < 0.01, *** p < 0.001, obtained with Mann-Whitney (HeLa cells, 24 h incubation time) or Student’s t-test.

As discussed above, weakly acidic uncouplers, acting as protonophores, enable a short-circuit for protons across the inner mitochondrial membrane. Thereby, the proton motive force (pmf) which is formed by an interplay of the mitochondrial membrane potential (Δψ_m_) and a proton gradient (ΔpH) across the membrane that drives the F_o_F_1_-ATPase, is dissipated. Thus, uncouplers inhibit the biosynthesis of ATP from ADP and inorganic phosphate (17, 26). Consequently, to restore the pmf, affected cells increase respiratory chain activity. This comprises enhanced substrate flux and increased oxygen consumption. Below a certain Δψ_m_ threshold, the F_o_F_1_-ATPase reaction is reversed from ADP phosphorylation to ATPase activity, which further depletes the ATP pool of affected cells (26).

The oxygen consumption and proton efflux rates (OCR and PER, respectively) of AETX-treated HeLa cells and fibroblasts were determined using a Seahorse XFe96 analyzer (Fig. S14). The Δψ_m_ was determined using the ratiometric fluorescent dye JC-10 in a plate reader-based assay. Both cell lines have been used in the past to compare effects on mitochondria (27). With sufficient glucose supply, like in our experiment, HeLa cells mainly rely on anaerobic glycolysis for ATP synthesis (28), while fibroblasts are known to largely rely on oxidative phosphorylation (29).

First, OCR and PER were measured directly after the addition of AETX to the cells, followed by the addition of oligomycin, 2-[[4-(trifluoromethoxy)phenyl]hydrazinylidene]propanedinitrile (FCCP) and rotenone/antimycin A (RAA). In both cell lines, the effects of AETX were dose-dependent throughout the experiments. Both cell lines showed an increase of OCR after acute injection of AETX (Fig. 4A, 4B, data on PER see Fig. S15 – S20). Although the magnitude of the effect was different, subsequent injection of oligomycin led to a decreased OCR and an increased PER in both cell lines, and the effect of FCCP on OCR and PER was diminished dose-dependently after AETX injection. Calculations according to Divakaruni *et al*. (26) revealed that both cell lines showed a significantly increased proton leak (Fig. S21) and decreased coupling efficiency (Fig. S22) compared to the control after treatment with AETX. These findings are indicative of uncoupling activity (26). To exclude any other cellular process apart from mitochondrial respiration being responsible for the observed increased oxygen consumption after AETX injection, we first inhibited respiratory chain activity with RAA before adding AETX to the cells (Fig. S23). We found no change in OCR, which is typical for uncouplers (17).

**Figure 4.**
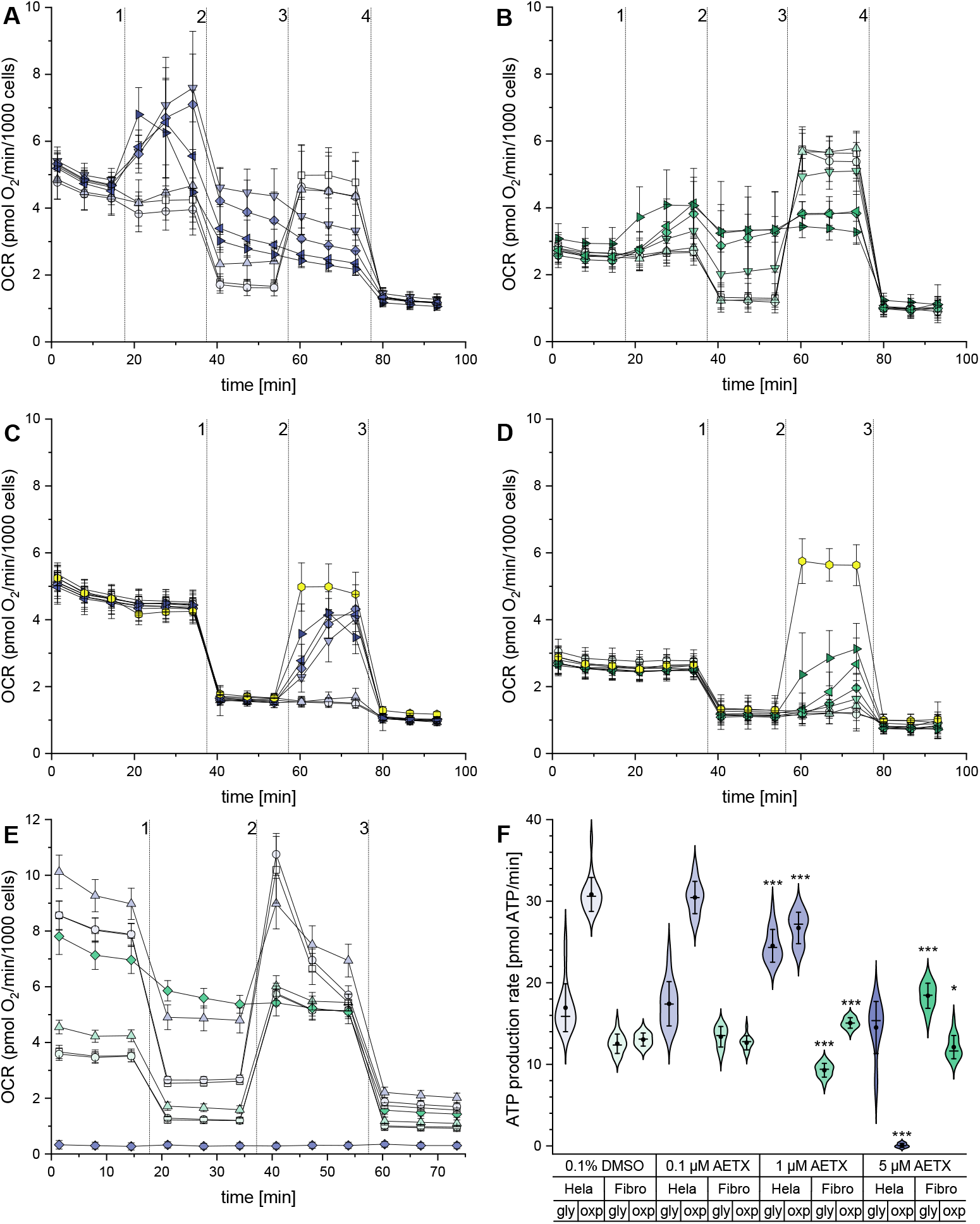
Influence of AETX on oxygen consumption and ATP production rates. Panels **A – F**: violet: HeLa cells, green: fibroblasts. Increasing shades of colour indicate increasing concentrations of AETX. Standard deviation (1x) indicated as whiskers. Panels **A – E**: □: control (0.1% DMSO), ○: 0.1 μM AETX, △: 1 μM AETX, ▽: 2 μM AETX, ◊: 5 μM AETX. Panels **A/B**: ◃: 10 μM, ▹: 30 μM AETX. Panels **C/D**: yellow ⬡: FCCP. Oxygen consumption rate measurement of **(A)** HeLa cells or **(B)** fibroblasts after acute stimulation with AETX. Addition of 1: AETX, 2: oligomycin, 3: FCCP, 4: rotenone/antimycin A + Hoechst 33342. Oxygen consumption rate measurement of **(C)** HeLa cells or **(D)** fibroblasts after acute stimulation with AETX or FCCP subsequent to FoF1-ATPase inhibition. Addition of 1: oligomycin, 2: AETX or 2 μM FCCP, 3: rotenone/antimycin A + Hoechst 33342. **(E)** Oxygen consumption rate measurement of HeLa cells and fibroblasts after 24 h of treatment with AETX. Addition of 1: oligomycin, 2: FCCP, 3: rotenone/antimycin A + Hoechst 33342. **(F)** ATP production rate in HeLa cells and fibroblasts respective to the origin of ATP. Statistically significant difference to the control indicated with * p < 0.05 and *** p < 0.001, obtained with Mann-Whitney (HeLa cells) or Student’s t-test (Fibroblasts). Mean indicated as black dot and the median as line.

Addition of AETX after inhibition of the F_o_F_1_-ATPase with oligomycin revealed a dose-dependent increase of OCR in both cell lines (Fig. 4C, 4D), indicating an increased respiratory chain activity, which again agrees with an uncoupling effect of AETX (26). In comparison to FCCP, the rise in OCR induced by AETX was lower. This could be explained by the fact that we titrated FCCP in preliminary experiments to obtain the optimal response from each cell line, since uncouplers are known to have a bell-shaped effect curve on OCR (30), and that we did not test the respective optimal concentration of AETX.

The effects of rotenone and antimycin A, both added after AETX to the cells, were not influenced by any AETX treatment, but were similar to the control throughout the experiments. This suggests that the targets of these compounds, complexes I and III of the respiratory chain, are not affected by AETX.

Uncoupling of the oxidative phosphorylation impacts the mitochondrial ATP production rate and interferes with the general energetic status of the cell (31–34). Therefore, we investigated the bioenergetic status of HeLa cells and fibroblasts after long-term incubation with AETX (Fig. 4E, 4F). Both cell lines were incubated for 24 h with 0.1, 1, and 5 μM AETX before measuring the OCR and PER, followed by calculations of the ATP production rates according to Desousa *et al*. (34). HeLa cells and fibroblasts treated with 0.1 μM AETX showed similar results compared to the control. At 1 μM AETX, the ATP production rate from anaerobic glycolysis in HeLa cells was enhanced, while ATP generated from oxidative phosphorylation decreased. However, in total, the ATP production rate increased compared to the control. A comparable effect was observed for fibroblasts after treatment with 5 μM AETX. Treatment of HeLa cells with 5 μM AETX led to a decrease of total ATP production, due to diminished oxidative phosphorylation. In contrast to treatment with 1 μM AETX, HeLa cells treated with 5 μM AETX did not adjust their glycolytic ATP production rate. This result agrees with our previous findings in the assays for cytotoxicity, as it might indicate that HeLa cells had already undergone cell cycle arrest, which is thought to be associated with lowered energy demand (35). However, affecting the electron transfer system may change the cytosolic NAD^+^/NADH+H^+^ ratio, which would alter glycolytic flux kinetics independently of cellular energy demand (32). Also, limitations in substrate transport due to diminished driving forces could contribute to this effect (26, 30). On the other hand, fibroblasts that were incubated with 1 μM AETX showed an increase of ATP from oxidative phosphorylation, while glycolytic ATP generation was decreased. As we could show that fibroblasts have a high spare respiratory capacity (Fig. S24), it is plausible that these cells could productively use the stimulation with 1 μM AETX and regulate ATP production accordingly.

To orthogonally assess the interference of AETX with mitochondrial ATP generation, we hypothesized that the cytotoxicity of AETX should be exacerbated in cells only capable of oxidative phosphorylation. In order to test this hypothesis, HCT116 cells were cultured in medium either containing a high glucose concentration to force cells into anaerobic glycolysis, or supplied only with pyruvate to force cells into oxidative phosphorylation. Indeed, we found an earlier onset of impairment (starting with 1 μM instead of 5 μM) and enhanced toxicity at 5 and 10 μM AETX in HCT116 cells cultivated in solely pyruvate containing medium when compared to the control and cells incubated in glucose containing medium (Fig. S25). This finding confirmed that AETX enhances the cellular need of glucose as energy source.

Having assessed oxygen consumption and ATP production rates, we strove to elucidate the effect of AETX on the Δψ_m_ in both cell lines (Fig. 3C, 3D). The Δψ_m_ was reduced in HeLa cells in all tested concentrations after acute stimulus and 2 h of incubation with AETX when compared to control. In fibroblasts, the Δψ_m_ rose when stimulated acutely with 5 and 10 μM AETX, increasing even further after 2 h of incubation, while cells treated with 0.1 and 1 μM displayed a reduced Δψ_m_. Again, the observed differences between both cell lines in reaction to AETX can be attributed to the lower maximum respiratory capacity of HeLa cells compared to fibroblasts. The maximum respiratory capacity is utilized to strengthen the Δψ_m_ consuming NADH-linked substrates, such as pyruvate, malate, or succinate (36). After 24 h of treatment, however, it can be assumed that substrate availability is reduced. Thus, the fibroblasts might no longer be able to fuel the increased respiratory chain activity, and therefore the Δψ_m_ decreases accordingly. Indeed, after 24 h of treatment, all concentrations of AETX resulted in a reduced Δψ_m_ in fibroblasts. This effect, determined in both cell lines, is characteristic for uncouplers (33, 36). In addition, a low Δψ_m_ is associated with cell cycle arrest in early G_1_-phase (35), which we indeed observed as described above.

Furthermore, the formation of reactive oxygen species (ROS) can be influenced by the interference of uncouplers with the pmf (37, 38). We deliberately chose 2′,7′-dichlorfluorescein-diacetate (DCF-DA) as a fluorescent probe to simultaneously assess cellular unspecific ROS level and mobilization of cytochrome c from complex VI of the respiratory chain as an indication of mitochondrial damage (39). We detected only a slight reduction of the DCF-DA signal after incubation of HeLa cells and fibroblasts with AETX (Fig. S26), suggesting that complex IV of the respiratory chain remains intact after treatment with AETX, and that ROS formation is largely unaffected in the first two hours after AETX treatment, but might be slightly reduced after 24 h of incubation.

### AETX acts as a protonophore on artificial lipid bilayers

In order to assess the capability of AETX to translocate protons across membranes in a protein-independent manner, we measured the conductance of a protein-free, artificial planar lipid bilayer separating two chambers containing aqueous solutions of different pH values in the absence or presence of AETX or FCCP as positive control. After adding hydrochloric acid to one of the chambers to obtain an aqueous solution of pH 3, we were able to detect electric currents in a voltage-dependent manner for FCCP as well as AETX (Fig. 5), demonstrating that AETX can shuttle protons across membranes. However, it has been reported that uncouplers might also interact with mitochondrial proteins to exert their effect on oxidative phosphorylation. Kotova *et al*. discussed that the effective concentration of certain uncouplers differs when tested on artificial lipid bilayers or in intact mitochondria. They concluded that active interaction of uncouplers with proton pumps or other proteins of the inner mitochondrial membrane might occur (30). Indeed, it has been shown that, e.g., FCCP or DNP increase the proton permeability of the mitochondrial membrane *via* induction of ADP/ATP carrier and uncoupling protein 1 activity (40). Our study is limited in this regard, as we can yet neither exclude nor suggest any additional interaction of AETX with mitochondrial proteins. However, the artificial lipid bilayer membrane assay demonstrated the capability of AETX to transport protons across membranes, confirming the underlying mechanism of action of AETX as a weakly acidic uncoupler.

**Figure 5.**
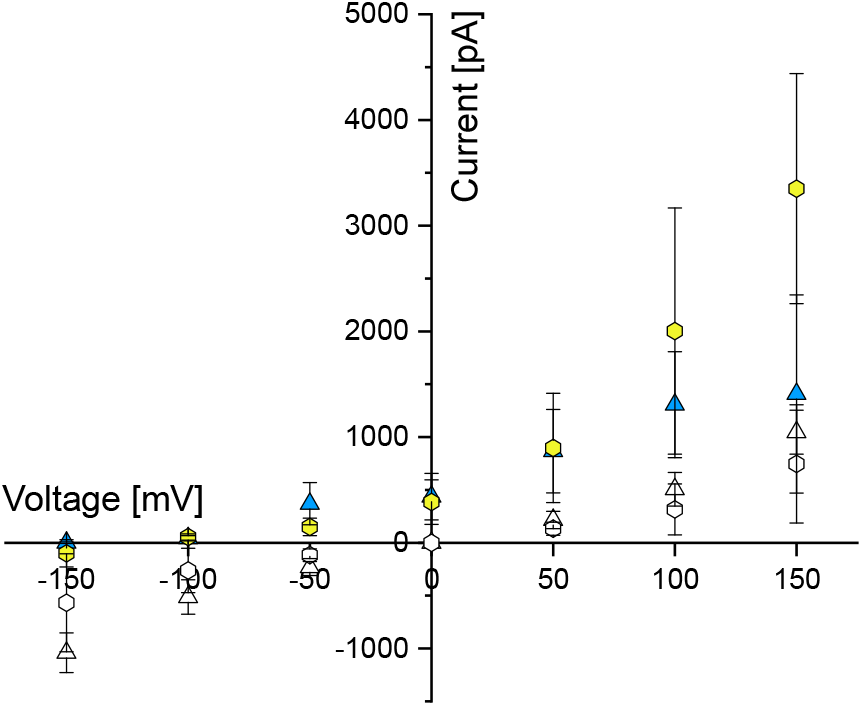
Current-voltage plot displaying translocation of protons across a protein-free artificial planar lipid bilayer. △: 1 μM AETX, ⬡: 1 μM FCCP. Yellow/blue: measurement in water with pH 3 on the amplifier side. White: control measurement in HEPES-buffered solution at pH 7. Whiskers indicate standard deviation (1x).

If AETX is indeed a weakly acidic uncoupler, its slightly acidic indole nitrogen should be crucial for its function as a protonophore, as it is known that blocking the acidic group of uncouplers, e.g., by methylation, abolishes their activity on the respiratory chain (41). We further hypothesized that removal of the electron withdrawing nitrile group should diminish its activity, since it should decrease NH acidity. To test these hypotheses, we synthesized *N*-methyl-AETX (m-AETX) by methylation of AETX using dimethyl sulfate. Desnitrile-AETX (dn-AETX) was isolated from the AETX-producing cyanobacterium *A. hydrillicola* as described before.^1^

Comparing the cytotoxicity of AETX and its two derivatives showed that after 24 h of incubation, m-AETX indeed was inactive at all tested concentrations (up to 10 μM), while dn-AETX, as expected, was less cytotoxic than AETX (Fig. 6A). The importance of the secondary amine and the nitrile group for the function of AETX as protonophore and, hence, uncoupler also became obvious when we compared the acute dissipation of the Δψ_m_ in HeLa cells induced by AETX, m-AETX and dn-AETX: While treatment with AETX led to immediate and concentration-dependent dissipation of the Δψ_m_, both derivatives rather hyperpolarized the mitochondrial membrane independent of the treatment concentration (Fig. 6B).

**Figure 6.**
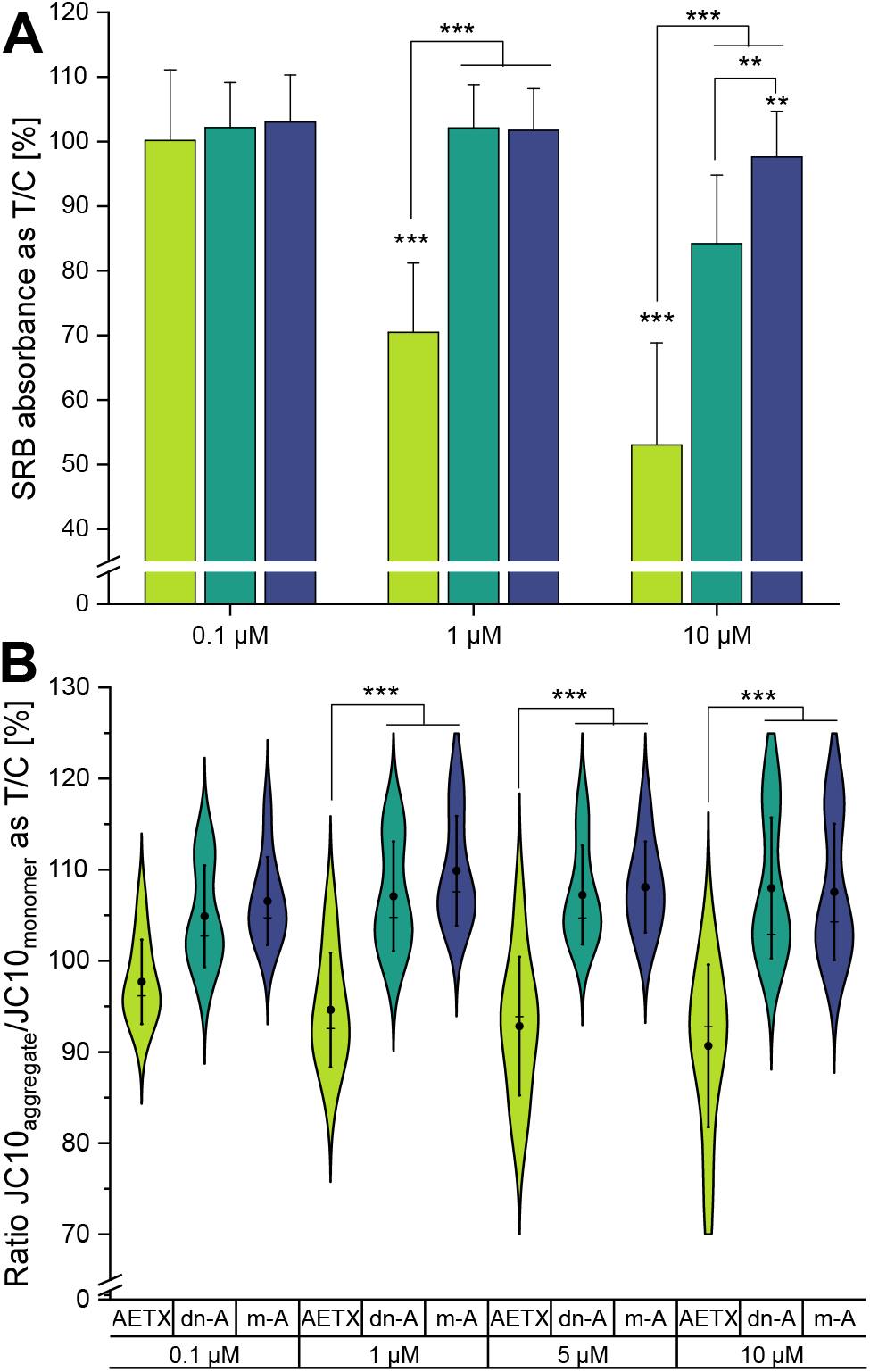
Comparison of the effect of AETX, *N*-methyl-AETX (m-AETX, m-A), and desnitrile-AETX (dn-AETX, dn-A) on overall cell viability and mitochondrial membrane potential. Panels **A/B:** light green: AETX, green: dn-AETX, blue: m-AETX. Standard deviation (1x) indicated as whiskers. **(A)** Cytotoxicity assay based on protein content (sulforhodamine B absorbance) in HCT116 cells normalized to the control. Statistically significant difference to the control indicated with ** p < 0.01 and *** p < 0.001, obtained with Mann-Whitney test. **(B)** Mitochondrial membrane potential as ratio of JC10 aggregate to JC10 monomer in treated HeLa cells normalized to the control. Mean indicated as black dot and the median as line. Statistically significant difference to treatment with AETX indicated with *** p < 0.001, obtained with Student’s t-test.

## Conclusion

We found that AETX acts as an uncoupler of oxidative phosphorylation. We assume that the cyto- and bacteriostatic effects we initially observed are due to a lack of ATP in the affected cells. This could also explain the observed morphological changes of *B. subtilis* from rod to spheric shape at concentrations below the MIC, as the elongation but not the division of *B. subtilis* is dependent on the ATP-binding cassette transporter-like complex FtsEX (42). For eukaryotic cells, perturbations in cellular energy homeostasis resulting in elevated AMP:ATP ratios are a known switch on the canonical AMP-dependent mechanism of AMP-activated protein kinase (AMPK) activation. Amongst other downstream effects, activated AMPK leads to enhanced mitophagy and cell cycle arrest in G_1_-phase (43). The reduced ATP production rate, the reduced TOM20 signals of mitochondria, and the cell cycle arrest in the G_1_-phase we observed in our assays suggest that AMPK activation is a downstream effect of AETX-induced uncoupling. Furthermore, AMPK activation in neurons induces Kv2.1 channel opening (44). Prolonged imbalances in neuronal ion homeostasis altering osmotic conditions in the brain could lead to the formation of the large intra-myelinic vacuoles observed in VM (45, 46). In addition, deficiency of the respiratory chain has been linked to myelin splitting at the minor dense lines like observed in animals suffering from AETX-induced VM (2, 47). However, these considerations need to be examined in more detail in future studies. The efficacy of AETX shown in the diversity of models used in our study is reflected by the broad range of affected animals, i.e., comprising taxa with and without myelin, emphasizing mitochondria as the universal primary target of AETX intoxication.

## Material and Methods Summary

Full experimental details and additional references for the methods can be found in SI. **Biology**. Bacteria were cultivated on LB-agar plates at 37 °C. The MIC was determined by measuring the OD_620_ of a bacteria suspension incubated for 20 h with AETX. Subcultures without any visible growth after 24 h were streaked on LB-agar plates and incubated for further 24 h to determine the minimal bactericidal concentration. Darkfield microscopy was used to visualize the morphology of *Bacillus subtilis*. Mammalian cell lines were cultivated in a humidified atmosphere at 37 °C and 5% CO_2_ in cell-line specific media. A sulforhodamine B assay was used to evaluate cytotoxicity. Confluence of AETX-treated HCT116 cells was monitored with time-course brightfield microscopy. To determine cell cycle arrest by flow cytometry, HCT116 cells were fixed in ice-cold ethanol and stained with RNase containing propidium iodide solution or FITC-conjugated Ki-67 antibody (Miltenyi Biotech). The Watson pragmatic model was used for cell cycle phase analysis. A formazan-based MTT-assay was used with PC-3-cells to determine a suitable concentration for the subsequent metabolomics analysis. After AETX treatment, PC-3 cells were fixed in an acidic ethanol solution and lysed by ultrasonication. After centrifugation, the supernatant was processed for LC-MS analysis (ion-pairing chromatography). Mass spectrometric analysis was conducted using targeted MS/MS with multiple reaction monitoring on a QTRAP 6500 system in neg. ion mode. Data processing and analysis was performed with the software MultiQuant and MetaboAnalyst. For immunofluorescence microscopy, HCT116 cells were grown on Ibitreat slides for cell imaging. After treatment, fixation, permeabilization, and blocking, cells were incubated with anti-TOM20 mouse monoclonal antibody (Santa Cruz Biotechnology), washed and incubated with the secondary Alexa Fluor 647 donkey anti-mouse antibody (Invitrogen). After mounting the cells with DAPI-containing mounting medium, images were acquired with a Confocal LSM Zeiss 800 system. To assess oxygen consumption and proton efflux rates, confluent HeLa cells and fibroblasts were treated with components of the Seahorse XF Cell Mito Stress Test Kit (Agilent), and AETX. Oxygen consumption and medium acidification was monitored with a Seahorse XFe96 Analyzer (Agilent). Data was normalized to cell count. The JC-10 Mitochondrial Membrane Potential Assay Kit for microplates (Abcam) was used to monitor mitochondrial membrane potential in HeLa cells and fibroblasts seeded in black 96-well plates with transparent bottom. Fluorescence was recorded in phenol red-free DMEM at 490/525 nm and 540/590 nm (excitation/emission). Similarly, the dye DCF-DA was applied to investigate unspecific reactive oxygen species formation. Fluorescence was measured in phenol-red free DMEM at 480/520 nm (excitation/emission). 500 μM H_2_O_2_ served as positive control. Origin software (OriginLab Corporation) was used for statistical analysis. All biological data were obtained from at least three independent biological replicates. **Chemistry**. Both pK_a_ and logP of AETX were determined using an automated titrator system with an incorporated UV-Vis spectrometer to acquire the spectrometric and potentiometric data (SiriusT3, Pion Inc.). A lipophilicity profile (logD *vs*. pH) was calculated from the experimentally determined pK_a_ and logP. To assess protonophore activity of AETX, a bilayer consisting of 1,2-diphytanoyl-sn-glycero-phosphatidyl-choline was formed covering an aperture (diameter 100 μm) in a Teflon septum separating two chambers containing an aqueous phase. Ag-AgCl reference electrodes were used to detect ion currents across the lipid bilayer. AETX and desnitrile-AETX were isolated from *A. hydrillicola* extract. m-AETX was synthesized from AETX using dimethyl sulfate. MS data of the compounds was acquired on an Orbitrap Exploris 240 mass spectrometer coupled to a Vanquish Flex HPLC system (Thermo Fisher Scientific). NMR spectra were recorded in DMSO-*d*_*6*_ on a Bruker Avance700. Compounds for biological assays were quantified using HPLC coupled with an evaporative light scattering detector (1290 Infinity II, Agilent).

### SI

The SI contains a detailed discussion of the structure confirmation of m-AETX, all experimental details, additional figures (including HPLC-UV chromatograms of AETX and derivatives, MS data, and NMR spectra of m-AETX), and data on the metabolomics experiments.

## Supporting information

SI metabolomics data

## Acknowledgements

We thank K.-A. Nave, Max-Planck-Institute for Multidisciplinary Sciences, Göttingen, Germany for helpful discussion, and S. Al-Robaiy, Center for Basic Medical Research (ZMG), University Hospital Halle (Saale), Martin Luther University Halle-Wittenberg, Germany for providing the infrastructure for the oxygen consumption rate measurements, and the core facility Multimodal Imaging of the Faculty of Chemistry of the University of Vienna for support of microscopy-based experiments. This research received no external funding. The mass spectrometers used in this study were partially funded by the German Research Foundation (DFG; project numbers 467315902. E. Dittmann, and INST 271/388-1, T.H.J.N.).

## Footnotes

1 Note to the reviewers: The manuscript describing the isolation and structure elucidation of dn-AETX from the cyanobacterium *Aetokthonos hydrillicola* is in revision at the *Journal of Natural Products* and will properly be cited in the final version of this manuscript.

