## Supplementary material for "Aetokthonotoxin, the causative agent of vacuolar myelinopathy, uncouples oxidative phosphorylation due to protonophore activity": SI

**Table of Contents**

| | Page |
|---|---|
| **SUPPLEMENTARY TEXT** | |
| Structure confirmation of *N*-methyl-AETX (m-AETX) | 2 |
| **METHODS – Biology** | 3 |
| **METHODS – Chemistry** | 6 |
| **SUPPLEMENTARY REFERENCES** | 8 |
| **SUPPLEMENTARY FIGURES** | |
| Microbiology | 10 |
| Cell biology – cytotoxicity | 11 |
| Metabolomics | 19 |
| Chemistry – $pK_a$ and logP | 22 |
| Cell biology – Seahorse experiments | 25 |
| Cell biology – enhanced cytotoxicity | 31 |
| Cell biology – ROS | 32 |
| Chemistry – purity of AETX, dn-AETX, m-AETX | 33 |
| Chemistry – structure confirmation of m-AETX | 33 |

**SUPPLEMENTARY TEXT**

**Structure confirmation of *N*-methyl-AETX (m-AETX).** The structure of m-AETX, and thus the successful *N*-methylation of AETX, was confirmed by 1D and 2D NMR spectroscopy and high-resolution mass spectrometry experiments based on the structure elucidation of AETX conducted by Breinlinger *et al.* (1). HRMS analysis resulted in a $[M+H]^+$ ion at m/z 667.6651, from which the molecular formula $C_{18}H_9N_3{}^{79}Br_2{}^{81}Br_3$ was calculated (calc. 667.6652, Δ 0.1 ppm). The presence of five bromo substituents in the molecule was also obvious from the observed isotope pattern. In contrast to AETX, which can only be ionized efficiently in negative ionization mode, m-AETX showed poor ionization efficiency, with ion signals detected exclusively in positive mode. The $^1H$ spectrum of m-AETX showed five distinct signals with chemical shifts in the aromatic region corresponding to the five protons of the two indole subunits. The chemical shifts (δ, in ppm) of the protons, the integration of these signals, and the coupling constants (J, measured in Hz) were as follows: δ 8.06 (d, J= 1.8 Hz, 1H), C-4'; δ 7.96 ppm (d, J= 1.8 Hz, 1H), C-6'; δ 7.79 ppm (dd, J= 1.8 Hz, J= 0.5 Hz, 1H), C-4; δ 7.52 ppm (dd, J= 8.8 Hz, J= 1.9 Hz, 1H), C-6; δ 7.40 ppm (dd, J= 8.8 Hz, J= 0.5 Hz, 1H), C-7, agreeing well with the data of AETX (1). An additional proton signal in the spectrum indicated the presence of a methyl group at δ 3.77 (s, 3H). In addition to the previously reported long-range correlations in AETX (key HMBC correlations are shown in Fig. S28, two additional long-range $^3J_{CH}$ couplings were observed in the HMBC spectrum: a correlation between the methyl group (*N*-methyl) and C-7a', and a correlation between the methyl group and C-2'. Based on these data, the successful *N*-methylation of AETX was confirmed. Fig. S29 shows the HRMS spectrum, Fig. S30, Fig. S31 and Fig. S32 show the $^1H$, $^{13}C$-HMBC and $^{13}C$-HSQC NMR spectra of m-AETX at 700 MHz.

**METHODS – Biology**

**Microbiology.** *Escherichia coli* DH5α (DSM 6897) and *Bacillus subtilis* WT168 (DSM 23778) were cultivated on LB-agar plates (Sigma-Aldrich, USA) at 37 °C.

**Minimal inhibitory concentration.** The minimal inhibitory concentration (MIC) of AETX was determined using the broth microdilution method according to the guidelines of the Clinical and Laboratory Standards Institute (CLSI) (2). In short, AETX was dissolved in DMSO and diluted in BD-BBL Mueller-Hinton-broth (MHB) (cation-adjusted, BD, USA) to various concentrations of 0.01-30 µM and adjusted to 0.3% DMSO in a clear 96-well plate (Greiner Bio-one, Germany). The inoculum was produced by picking colonies, suspending them to an optical density ($OD_{600}$) of 0.08-0.10 and diluting 20-fold to inoculate a total of $5\times10^4$ CFU/well. After an incubation of 20 h with AETX at 37 °C, the bacterial growth was determined by measuring the OD at 620 nm using a Tecan Infinite 200 Pro M Plex plate reader (Tecan Group, Switzerland). An $OD_{600}$ of 0.02 higher than the uninoculated control was defined as growth. MICs were determined as the median of biological triplicates with five technical replicates each.

**Minimal bactericidal concentration.** The minimal bactericidal concentration (MBC) was investigated according to the CLSI guideline (3) by incubating the 96-well plates described above to a total of 24 h at 37 °C. Then, 10 µL subcultures which did not show any visible growth were streaked onto LB-agar plates. After another 24 h incubation period, the colonies were counted. The determination was performed in biological triplicates with two technical replicates each.

**Morphological changes of *B. subtilis*.** To visualize the effect of AETX on *B. subtilis* cells, an Olympus CX31 microscope with a PlanC N 40x/0.65 Ph2 objective (Olympus, Japan) was used in dark field mode after 24 h incubation for image acquisition. Pictures were taken with a MikrOcular Full HD ocular camera and analyzed using the MikroCamLabII software (both Bresser, Germany).

**Cell biology – cell lines, reagents and cultivation.** Human colorectal cancer HCT116 cells (ACC 581, DSMZ-German collection of Microorganisms and Cell Cultures, Germany), human cervix carcinoma HeLa cells (provided by Prof. Junker, Martin-Luther-University, Halle, Germany), human foreskin fibroblasts CCD1092Sk cells (immortalized cell line, provided by Prof. Gekle, Julius-Berstein-Institut, Halle, Germany), and human prostate cancer PC-3 cells (purchased from ATCC, USA) were maintained at 37 °C in humidified atmosphere with 5% $CO_2$. Cells were passaged after reaching 80% - 90% confluence. HeLa and CCD1092Sk cells were cultivated in Dulbecco's modified eagle medium (DMEM) with low glucose (ROTI® CELL DMEM Low Glucose, Carl Roth, Germany) completed with 10% (v/v) fetal bovine serum (FBS, Sigma Aldrich, USA) and 2 mM glutamine (ROTI®-CELL Glutamin-Lösung, Carl Roth, Germany), while HCT116 cells were maintained in McCoy's 5A medium (ROTI®Cell McCoy's 5A, Carl Roth, Germany) supplied with 10% (v/v) FBS, penicillin (100 I.U./mL) and streptomycin (100 mg/L) (Pen/Strep-PreMix, Carl Roth, Germany). PC-3 cells were cultured in RPMI 1640 medium, supplemented with 10% (v/v) FBS, 1% (v/v) L-glutamine, penicillin (100 I.U./mL) and streptomycin (100 mg/L) (media and supplements purchased from Capricorn Scientific, Germany), and used for a maximum of six passages per batch. Aetokthonotoxin (AETX) and derivatives (purity > 95%, Fig. S27) were dissolved in DMSO (AppliChem, Germany) and diluted in the respective media used in the individual experiments. For each assay, at least three independent biological replicates were performed (unless stated otherwise). The respective solvent control was always carried along during the course of each experiment.

**Sulforhodamine B assay.** The sulforhodamine B (SRB) colorimetric assay was conducted as previously described (4). Briefly, HeLa cells, fibroblasts or HCT116 cells were seeded on a clear, cell culture treated 96-well plate with flat bottom. The next day, cells were treated with 0.1, 1 and 10 µM of AETX, m-AETX or dn-AETX, respectively, and incubated for 24 h, fibroblasts were incubated for 48 h, due to their longer doubling time. After the respective incubation time, cells were directly fixed with cold 10% (w/v) trichloroacetic acid (TCA) for one hour. Then, cells were carefully rinsed four times with slow-running tap water and blow dried. When cells were completely dry, 0.057% (w/v) of SRB solution (Sigma Aldrich, USA) in 1% (v/v) acetic acid was added to the wells. After 30 min incubation time at room temperature, cells were quickly washed four times with 1% (v/v) acetic acid and blow dried. Finally, 10 mM Tris base solution (pH 10.5) was added to the completely dry wells, the plate was placed in a TECAN Infinite M Plex plate reader, orbitally shaken for 300 s, and absorbance was recorded at 510 nm. In addition to that, AETX was submitted to the National Cancer Institute Developmental Therapeutics Program. There, AETX was tested on 59 cancer cell lines under use of the SRB assay in a concentration range of 5 nM to 50 µM. To evaluate toxicity of AETX in dependence on substrate availability, we cultivated HCT116 cells in two different media: DMEM-glucose (DMEM high glucose (25 mM) + 10%

FBS + penicillin (100 I.U./mL) + streptomycin (100 mg/L) + 2 mM glutamine + 25 mM HEPES) and DMEM-pyruvate (DMEM no glucose + 5 mM pyruvate + 10% FBS + penicillin (100 I.U./mL) + streptomycin (100 mg/L) + 2 mM glutamine + 25 mM HEPES) for 24 h before treatment with AETX. AETX was diluted in the respective DMEM-glucose or DMEM-pyruvate medium, and cells were incubated for 24 h with 0.1% DMSO, 0.1, 1, 5 or 10 µM AETX. Then, we conducted the SRB assay as described above.

**Morphological changes of HeLa cells.** Phase contrast microscopy was used to observe morphological changes of AETX-treated HeLa cells. HeLa cells were seeded in transparent 96-well plates, allowed to settle for 24 h and then treated for further 24 h with 0.3% DMSO, 0.1, 1, 5, 10 and 30 µM AETX at 37 °C and 5% $CO_2$ in a humidified atmosphere. At the end of the incubation time, phase contrast images were acquired at room temperature and ambient air with an inverse microscope (Axio Observer, Zeiss, Germany), equipped with 10, 20, and 63x objective lenses, an Axiocam 712 color digital camera and ZEN 3.2 software (blue edition, Zeiss, Germany).

**Confluency determination.** To obtain a first impression on cytostatic activity of AETX, HCT116 cells were seeded in transparent 96-well plates. After 24 h, confluence was assessed per well using brightfield microscopy to calculate the confluency of the obtained images (Axion Lux microscope and CytoSmart software, both CytoSmart Axion Biosystems, USA). Then, cells were incubated with various concentrations of AETX and imaged again after 24 h and 48 h.

**Cell cycle arrest.** HCT116 cells were seeded in 12-well plates, allowed to attach overnight, and incubated for 24 h with various concentrations of AETX. Then, cells were detached from the plates, set to $1\times10^6$ cells per condition and fixed in ice-cold 70% ethanol. Fixed cells were stored at -20 °C until the day of the flow cytometry experiment. To determine the cell cycle phase, fixed cells were stained for 30 min with propidium iodide (PI) solution containing RNase (FxCycle PI/RNase, Thermo Fisher, USA) and analyzed with a CytoFlex flow cytometer (Beckman Coulter, USA) according to manufacturer's instructions. To discriminate between cells in $G_0$ and $G_1$ phase, fixed cells were stained with Ki-67 FITC conjugated antibody (Miltenyi Biotech, Germany) for 20 min, washed in PBS and then analyzed. The software CytExpert v.2.6 (Beckman Coulter, USA) and the free web-based software floreada.io (https://floreada.io) utilizing the Watson pragmatic model were used for gating and cell cycle phase analysis. $2\times10^4$ cells were analyzed per technical replicate. Three technical replicates per biological replicate were examined. Cells were gated in the following way: First, cell debris was excluded and cell singlets were selected. Then, a gate was set to separate autofluorescence using unstained cells and the appropriate channel (PE for propidium iodide, FITC for FITC-conjugated Ki-67 antibody).

**Metabolomics – MTT assay for treatment concentration determination.** Cells were seeded in 96-well plates at 6000 cells per well in 100 µL of medium and allowed to adhere overnight. The cells were then treated with AETX at eight concentrations in 0.5% DMSO. Controls included 0.5% DMSO (negative) and 100 µM digitonin (positive, set as 0% viability; Riedel De Haën, Germany). Each experiment comprised two biological replicates and four technical replicates. After 48 h, cells were washed with PBS and incubated with MTT solution (0.5 mg/mL; Sigma-Aldrich, Germany) for 1 h under standard growth conditions. Formazan dye, formed by the portion of viable cells, was subsequently dissolved with DMSO, and absorbance was measured at 570 nm, with 670 nm as a background reference using a SpectraMax M5 plate reader (Molecular Devices, USA). Cell viability was calculated relative to untreated controls. Data analysis and $IC_{50}$ calculations were performed using GraphPad Prism 10 and Microsoft Excel 2013, employing a four-parameter logistic function for mean values. $IC_{50}$ was determined to be 0.74 µM AETX (Fig. S9). This concentration was subsequently used for the metabolomic assay.

**Metabolomics – sample preparation.** Metabolomics samples were proceeded in accordance with the protocol outlined before (5). In brief, cells were seeded in T-25 flasks with six replicates for each condition. After adhering overnight, cells were treated with solvent (0.0074% DMSO), or AETX at $IC_{50}$ concentration determined from a previous viability assay ($IC_{50}$ in the MTT assay: 0.74 µM). After 48 h of incubation under standard growth conditions, untreated control cells reached 85% confluency. In addition, similar independent experiments were conducted using 2 h, 4 h, and 24 h incubation time with AETX. Subsequently, the cells were washed with pre-warmed PBS (37 °C). Subsequently, cells were rapidly fixed with cold acidic ethanol solution (10% v/v HCl, pH 1.4, -80 °C). Tightly sealed flasks were submerged in an ultrasonic water bath with dry ice to maintain a low temperature (around 4 °C) while undergoing 5 min of sonication to detach the cells. The cell suspension was transferred to pre-chilled Eppendorf tubes on dry ice. Sample volumes were reduced to 50 µL using a nitrogen stream. Two rounds of centrifugation were performed at 10600 rcf for 5 min at 1 °C, and the supernatants were transferred after each step to fresh tubes. The final supernatant was placed into LC-MS vials and stored at -80 °C until analysis.

**Metabolomics – analysis.** The metabolomics experiments were conducted following the protocol recently described (5). In brief, hydrophilic metabolites were separated using ion-pairing chromatography on a Nucleoshell RP18 column (2.1 × 150 mm, 2.1 µm particle size; Macherey & Nagel, Germany) with a Waters ACQUITY UPLC System, featuring an ACQUITY Binary Solvent Manager and Sample Manager (injection volume of 5 µL; Waters, Germany). The mobile phases were 10 mmol/L tributyl amine (pH 6.2, adjusted with glacial acetic acid) for eluent A and acetonitrile for eluent B. The elution profile started with 2% (v/v) of eluent B in A held isocratically for 2 min, followed by a linear gradient increasing from 2% to 36% (v/v) B in A over 16 min. It then continued from 18 to 21 min up to 95% (v/v) B in A, held isocratically from 21 to 22.5 min, and decreased back to 2% (v/v) B in A from 22.51 to 26 min. The flow rate was 400 µL/min, and the column temperature was maintained at 40 °C. Mass spectrometric analysis of the central carbon and cellular energy metabolism metabolites was conducted using targeted MS/MS with multiple reaction monitoring (MRM) on a QTRAP 6500 system (AB Sciex, Germany), operating in negative ion mode and managed via Analyst 1.7.1 software (AB Sciex, Germany). Key settings included: ion spray voltage at -4500 V, nebulizing gas at 60 psi, source temperature at 450 °C, drying gas at 70 psi, and curtain gas at 35 psi. Data processing involved peak integration using MultiQuant software version 3.0.3 (Sciex, USA). To correct for variations of the cell numbers between treatments and controls, the metabolite peak areas were normalized to the total peak area per sample. Each normalized metabolite signal was then divided by the mean of the normalized control signals within the same experiment. The resulting data were $\log_2$-transformed. Statistical analysis and data visualization were performed using MetaboAnalyst 6.0 software, employing $\log_2$-normalized data and Range Scaling for consistency.

**Immunofluorescence microscopy.** HCT116 cells were seeded on Ibitreat slides for cell imaging (Ibidi, Germany). Cells were treated for 24 h with 0.1, 1, and 5 µM AETX, washed with PBS, and fixed with 3.5% (w/v) formaldehyde solution for 15 min in the dark. Afterwards, cells were washed with PBS once, then with a PBS-glycine solution (0.375 g glycine in 50 ml PBS-A), followed by two more PBS washing steps. Subsequently, cells were permeabilized with 0.2% Triton-X in PBS, blocked with 2% donkey serum for 1 h, and incubated with primary anti-TOM20 antibody (diluted 1:500 Santa Cruz Biotechnology, USA) for 2 h. Then, cells were washed three times with 0.05% Triton-X in PBS and twice with PBS. Cells were incubated with the secondary antibody AlexaFluorTM 647 donkey anti-mouse IgG (Invitrogen, USA), diluted 1:1000, overnight. The previous washing steps were repeated, and cells were submerged in two drops of mounting medium containing DAPI (Roth, Germany). Images were acquired with a confocal LSM Zeiss 800 system using a Plan Apochromat63X/1.4 Oil DICII objective. Image analysis results from the quantification of n > 100 regions of interest of the mitochondrial network with ImageJ v. 1.53t. Every experimental condition was measured in quadruplicates.

**Oxygen consumption, proton efflux, and ATP production rate.** A Seahorse XFe96 Analyzer (Agilent, USA) was used to determine cellular oxygen consumption rate (OCR) and proton efflux rate (PER). $2\times10^4$ HeLa or CCD1092Sk cells were seeded per well in Seahorse XF cell culture microplates (Agilent, USA) and allowed to attach for 24 h. On the day of the experiment, medium was changed to Seahorse XF DMEM, pH 7.4, supplemented with 10 mM glucose, 2 mM glutamine and 1 µM pyruvate (all Agilent, USA). Brightfield images of the cells were taken at 37 °C and 0% $CO_2$ using a Cytation 1 imaging reader (BioTek) to check for confluence. Meanwhile, a Seahorse XF cell mito stress kit (Agilent, USA) was prepared according to the manufacturer's instructions, as were treatment concentrations of AETX. The hydrated sensor cartridge was equipped according to the experimental requirements with various concentrations of AETX or solvent control and the kit components. For HeLa cells, 2 µM oligomycin, 0.5 µM 2-[[4-(trifluoromethoxy)phenyl]hydrazinylidene]propanedinitrile (FCCP) and 0.5 µM rotenone + antimycin A (RAA), for CCD1092Sk cells, 2 µM oligomycin, 2 µM FCCP and 0.5 µM RAA were applied. The last compound addition was supplemented with 3 µL/mL Hoechst 33342. These concentrations and seeding densities were determined as optimal in preliminary experiments. After one hour incubation at 37 °C without $CO_2$, the plate was placed into the Seahorse XFe96 Analyzer to assess OCR and PER at 37 °C. Each measurement (baseline or after compound addition) consisted of 3 min mixing and 3 min measuring and was repeated three times. After the last measurement, fluorescence images were taken with the imaging reader for cell number calculation. OCR and PER were normalized to the number of cells per well. For the 24 h experiment, cells were incubated for 24 h with three concentrations of AETX. On the following day, the experiment was conducted the same way as described above. The software Wave v. 2.6.3.5 and Agilent Technologies Cell imaging v. 1.1.0.17 (both Agilent, USA) were used for data analysis. Coupling efficiency was calculated according to (6) and the

Agilent user guide RA.4773611111. ATP production rate from glycolysis or oxidative phosphorylation was evaluated using the equations for calculation of ATP production rates recently presented in (7) using PER instead of extra cellular acidification rate (ECAR). Fig. S14 provides a supporting sketch for the interpretation of the results.

**Mitochondrial membrane potential.** To assess the mitochondrial membrane potential, HeLa and CCD1092Sk cells were seeded in a black 96-well plate with flat, clear bottom and allowed to attach overnight. Cells were then incubated either for 2 h or 24 h with various AETX concentrations. Alternatively, to capture and compare immediate effects, mitochondrial membrane potential was measured instantly after addition of AETX or its derivatives (m-AETX) and (dn-AETX) to the cells. The JC-10 Mitochondrial Membrane Potential Assay Kit for microplates (Abcam, UK) was used following the manufacturer's instructions. Fluorescence of JC-10 was recorded with a TECAN Infinite M Plex plate reader at excitation/emission 490/525 nm and 540/590 nm.

**Reactive oxygen species.** The general reactive oxygen species (ROS) status was assessed in HeLa and CCD1092Sk cells using the fluorescent dye DCF-DA (Sigma Aldrich, USA). Cells were seeded in black 96-well plates with clear flat bottom (Corning, USA) and allowed to attach overnight. The following day, cells were incubated for 24 h with various concentrations of AETX to assess the long-term effect of AETX. Then, cells were washed and incubated for 30 min with 50 µM DCF-DA in phenol red-free DMEM (PAN Biotech, Germany). After washing the cells twice, fluorescence was measured in phenol red-free DMEM at excitation/emission 480/520 nm in a Tecan Infinite M Plex plate reader. For the short-term kinetic measurement, on the day after seeding cells were directly incubated for 30 min with 50 µM DCF-DA as stated above. After the washing steps, background fluorescence was measured before AETX in various concentrations and 500 µM $H_2O_2$ (positive control) were added to the cells. Fluorescence was recorded directly and 15, 30, 60 and 120 min after treatment, using the same setting as mentioned above.

**Statistic.** Origin software v. 9.6.0.172 (OriginLab Corporation, USA) was used for statistical analysis. First, normality was assessed with Lillie-Force test for normality. Then, parametric data was assessed using two-sided student's t-test, while non-parametric data was assessed using Mann-Whitney test. Sigmoidal fit was calculated using the same software.

**METHODS – Chemistry**

**p$K_a$ determination.** An automated titrator system with an incorporated UV-Vis spectrometer (SiriusT3, Pion Inc., USA) was used to acquire the spectrometric data.The optical system consisted of a photodiode array detector with a deuterium lamp and a fibre optic dip probe, the titrator module comprised a temperature controller (by Peltier device with *in-situ* thermocouple), a pH electrode, an overhead stirrer, and motorized dispensers for the automatic delivery of assay titrants and reagents via capillaries. The instrumentation was operated using SiriusT3Control software (V2.0). All experiments were carried out at a controlled temperature 25.0 ± 0.2 °C. The pH range of titration assays was set between pH 2.0 to pH 12.0. Prior to use, 0.5 M KOH base titrant was standardized by the titration of approximately 15 mg of potassium hydrogen phthalate, in triplicate. 0.5 M HCl titrant was subsequently standardized against the base titrant. The assay media for p$K_a$ determination was kept at a constant ionic strength of 0.15 M KCl and under argon atmosphere. The pH electrode was calibrated daily using the Avdeef-Bucher four-parameter equation (8). HPLC grade methanol cosolvent was used. The aqueous p$K_a$ was determined by Yasuda-Shedlovsky extrapolation (9) from mixtures of water and methanol. Data processing and generation of the reported p$K_a$ values was carried out using SiriusT3Refine software (V2.0).

**logP determination.** An automated titrator system (SiriusT3, Pion Inc., USA) was used to acquire the potentiometric data. The aqueous p$K_a$ value of the sample was pre-measured for the calculation of logP from the potentiometric data. Ionic strength adjusted (0.15 M KCl) water and 1-octanol (pre-saturated with ionic strength adjusted water) were added to 0.6 - 0.8 mg AETX. The p$K_a$ in water (pre-measured aqueous p$K_a$) and the apparent p$K_a$ in the presence of octanol (p$_o$$K_a$), were compared and the logP was determined (10). Using the experimentally determined p$K_a$ and logP, a lipophilicity profile (logD *vs.* pH) of the compound was calculated. The potentiometric method for the determination of logP has been thoroughly validated (11, 12).

**Planar lipid bilayer assay.** Planar lipid bilayers were formed as described before (13, 14). Briefly, two home-made black Delrine half-cuvettes of 2.5 mL were used to sandwich a Teflon septum (20 µm thickness, Goodfellow, GB) having an aperture with a diameter of 100 µm. The surrounding was prepainted with hexadecane dissolved in n-hexane at 1-5% (v/v), and the compartments (2.5 mL) were dried for 30-35 min in order to evaporate the solvent. Standard Ag-AgCl reference electrodes with a diaphragm (Metrohm, Germany) were used to detect the ionic current. One electrode was grounded, whereas the other was linked to the headstage of an Axopatch 200B amplifier (Axon Instruments from Molecular Devices, USA), used for the conductance measurements in voltage clamp mode. The signals were filtered by an on-board low pass Bessel filter at 1 kHz and with a sampling frequency of 10 kHz. Examination of the current recordings was completed using Clampfit (Axon Instruments). The current-voltage relation of the individual experiments was determined from single averaged currents at given voltages. The experiment was performed in a temperature-regulated (20 °C) and electrically screened room. The bilayers were made by adding 10 µL of 1,2- diphytanoyl-sn-glycero-phosphatidyl-choline (Avanti Polar Lipids, USA) at a concentration of 5 mg/mL in n-pentane on top of each half-cuvette (area about 1 $cm^2$ each). The solution was buffered with 10 mM HEPES (Sigma Aldrich, USA) at pH 7. We first measured the conductance of the bilayer membrane alone, which was negligible. After ensuring a tight membrane, we added 1 µM of AETX or FCCP, both dissolved in DMSO, to the aqueous phase. Due to the geometric constraint having a micrometer-sized lipid patch on one side of the cuvette, equilibrium partitioning of AETX or FCCP is difficult to achieve. Compound addition to the lipid phase caused instability of the membranes. To accelerate equilibrium, we repeatedly broke and reformed the lipid bilayer. At lower concentrations, equilibrium was difficult to achieve, whereas higher concentrations resulted in an instable lipid membrane. To test proton transport we exchanged buffered solution on the amplifier side and replaced the solution with water. We titrated in several steps HCl and measured the ion current to perform an I-V plot. Smaller pH gradients showed similar trends but with high fluctuations (data not shown).

**Purification of AETX and desnitrile-AETX.** Aetokthonotoxin (AETX) and desnitrile-AETX (dn-AETX) were isolated from *A. hydrillicola* extract as described before.[1] AETX and dn-AETX containing biomass was harvested and lyophilized. The biomass was suspended in 50% MeOH (v/v), homogenized by vortexing, treated with an ultrasonic rod (Bandelin, Germany) and extracted on a shaker for 20 min. After centrifugation, the biomass pellet was subsequently extracted again in the same manner with 50% MeOH (v/v), and twice with 80% MeOH (v/v). All supernatants were combined and dried *in vacuo*. The extract was fractionated using flash chromatography on a $C_{18}$ cartridge (CHROMABOND® Flash RS 80 $C_{18}$ec, 15-40 µm, 30.9 × 49 mm, Macherey Nagel, Germany) on a preparative HPLC System (Gilson, USA). A binary gradient from 30-60% MeOH (v/v) in water in 6 min, 60-100% MeOH (v/v) in water for a further 18 min, and 100% MeOH for 10 min has been used. Fractions were collected every 2 min and dried in a vacuum centrifuge. The fractions containing AEXT and dn-AETX were redissolved in 2 mL of MeCN 80 (v/v) and subjected to semi-preparative HPLC (Dionex UltiMate 3000, Thermo Fisher, USA) using a Luna PFP2 column (250x10 mm, 5 µm, 100 Å, Phenomenex, USA). The following chromatographic parameters were used: binary gradient from 64-98% (v/v) MeCN in water (0.1% formic acid each) at 5 mL/min in 17 min. For purity control the following chromatographic parameters were used: Kinetex C18 column (50 × 2.1 mm, 2.6 µm, 100 Å, Phenomenex, USA), binary gradient from 5-100% (v/v) MeCN in $H_2O$ (0.1% formic acid each) at 0.4 mL/min in 18 min, 100% MeCN for 2 min, performed on a 1290 Infinity II (Agilent, USA). Purity has been confirmed to be >99.5%.

**HPLC-HRMS data acquisition.** HRMS data was acquired on an Orbitrap Exploris 240 mass spectrometer (Thermo Fisher Scientific, USA) equipped with a heated ESI interface coupled to a Vanquish Flex HPLC system (Thermo Fisher Scientific, USA). Chromatographic parameters: Kinetex $C_{18}$ column (50 × 2.1 mm, 2.6 µm, 100 Å, Phenomenex, USA), binary gradient from 5-100% (v/v) MeCN in water (0.1% formic acid each) at 0.4 mL/min in 16 min, 100% MeCN in 4 min. HRMS data acquisition was conducted in positive and negative ionization mode, ESI spray voltage 3.5 kV and -2.5 kV, capillary temperature 300 °C, sheath gas flow rate 40 L/min, auxiliary gas flow rate 5 L/min. Full scan accurate mass spectra were acquired from *m/z* 133.4 to 2000 with a resolution of 35 000 at *m/z* 200.

**Methylation of AETX.** AETX was dissolved in THF. Under stirring at room temperature, dimethyl sulfate (3.0 equiv.) and potassium carbonate (5.0 equiv.) were added. After stirring for 30 min at room temperature, the mixture was concentrated under reduced pressure and purified by HPLC. $R_f$ (toluene/hexane 2:1) = 0.7.

**NMR data acquisition.** NMR spectra were recorded in DMSO-$d_6$ on a Bruker Avance700 (Bruker BioSpin, Germany) operating at 700 MHz ($^1H$) or 175 MHz ($^{13}C$) at 300 K. Chemical shifts in ppm relative

[1] Note to the reviewers: The manuscript describing the isolation and structure elucidation of dn-AETX from the cyanobacterium *Aetokthonos hydrillicola* has been submitted to the *Journal of Natural Products* and will properly be cited in the final version of the manuscript."

to the residual solvent chemical shifts ($\delta_H$ 2.50, $\delta_C$ 39.52). NMR data were analyzed with MestReNova (version 14.3.0-30573, Mestrelab Research, Spain) after Auto Phase Correction and Auto Baseline Correction.

**Quantification of Desnitrile-AETX (dn-AETX) and *N*-methyl-AETX (m-AETX).** The concentrations of test compound solutions for bioactivity testing were quantified using HPLC coupled with an evaporative light scattering detector (1290 Infinity II, Agilent, USA) as described previously (15). Synthetic AETX (16) was used as standard to establish a calibration curve from 23.7 to 237 ng on-column. 1, 2.5, 5 and 10 µL of a 23.7 ng/µL solution in 90% (v/v) MeCN in $H_2O$ were injected in triplicate on a Kinetex C18 column (100 × 3 mm, 2.6 µm, 100 Å, Phenomenex), and eluted with a gradient from 10-100% (v/v) MeCN in $H_2O$ (0.1% FA each) over 10 min at 0.65 mL/min. Settings of the ELSD were as follows: evaporator temperature 45 °C, nebulizer temperature 45 °C, gas flow rate 1.3 standard liter per minute, $N_2$ 3.5 bar. The calibration curve was generated as described by Young et al. (17). In brief, the response areas were averaged, and log(ELSD response area) was plotted against log(amount in ng) to generate a linear calibration curve. dn-AETX and m-AETX were dissolved in 1 mL MeCN 90% (v/v in dd$H_2O$), diluted 1:3 in the same solvent, and injected in triplicate under the same conditions.

**SUPPLEMENTARY FIGURES**

**Microbiology**

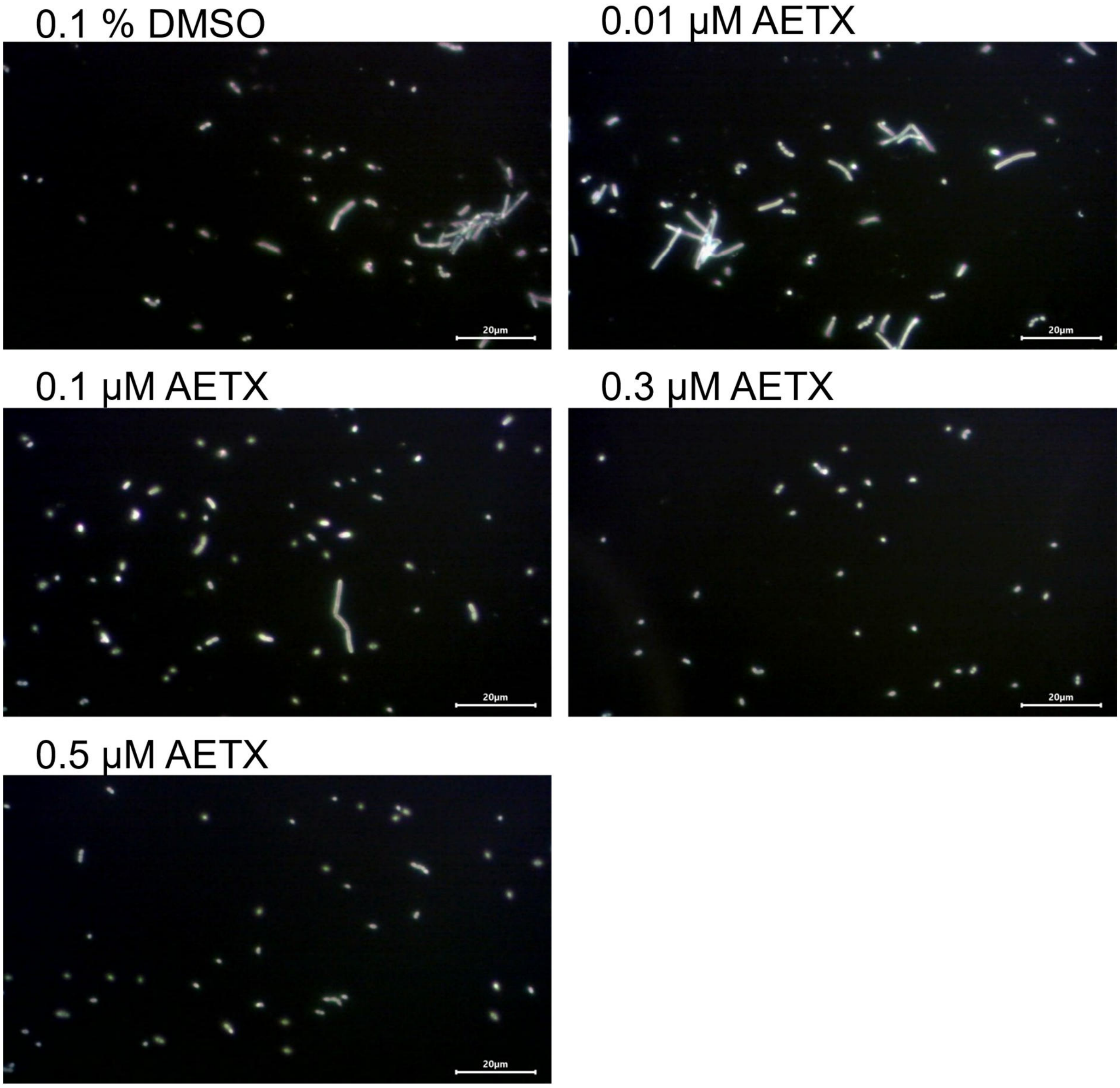

**Fig. S1:** Darkfield microscopy images of morphological changes from rod to spheric shape of *B. subtilis* after 24 h incubation with 0.1% DMSO or 0.01, 0.1, 0.3, and 0.5 µM AETX. Scale bar 20 µm.

### Cell biology – cytotoxicity

**A**

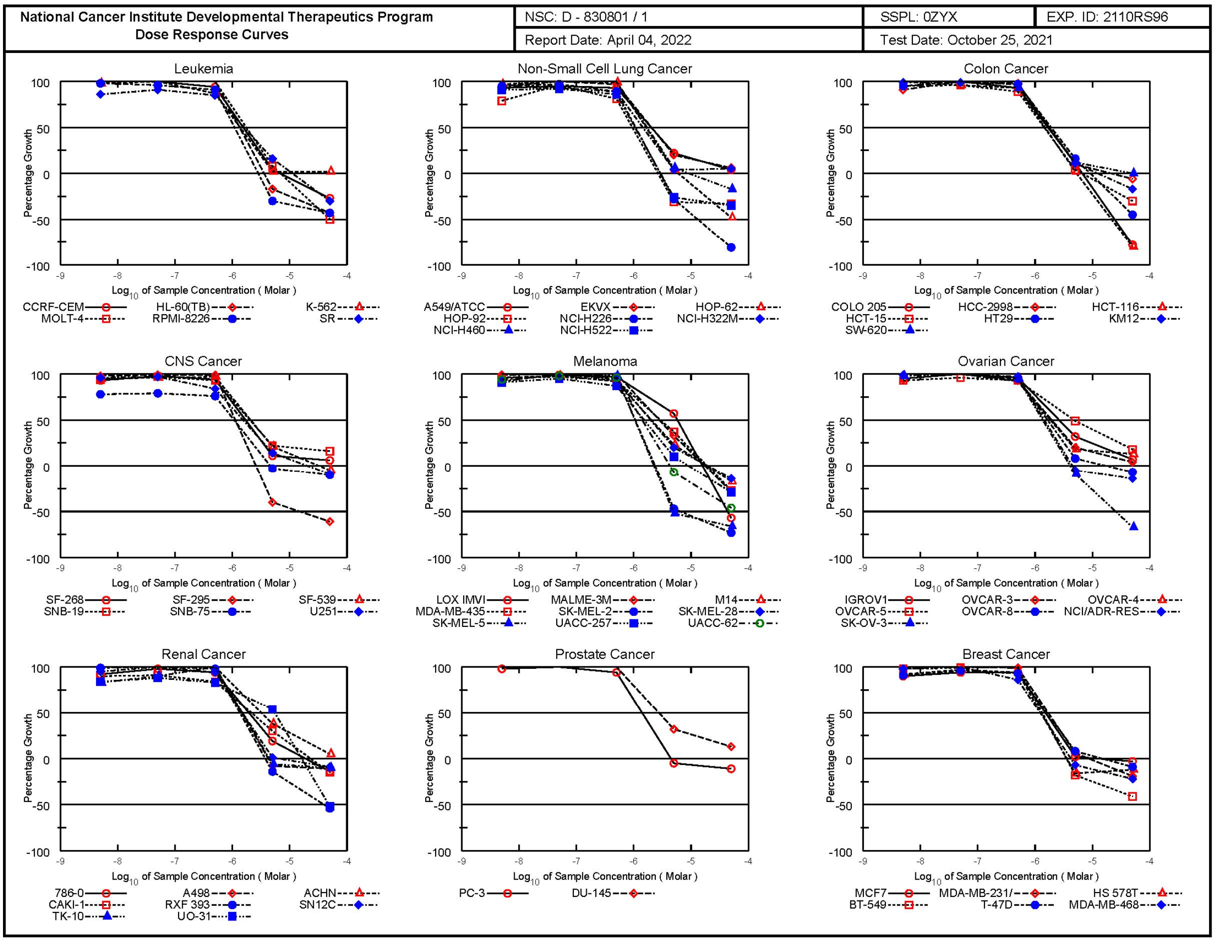

**B**

### National Cancer Institute Developmental Therapeutics Program
### In-Vitro Testing Results

| | | | |
|---|---|---|---|
| NSC : D - 830801 / 1 | Experiment ID : 2110RS96 | Test Type : 08 | Units : Molar |
| Report Date : April 04, 2022 | Test Date : October 25, 2021 | QNS : | MC : |
| COMI : AETX | Stain Reagent : SRB Dual-Pass Related | SSPL : 0ZYX | |

| | Time | | Log10 Concentration | | | | | | | | | | | | |
|---|---|---|---|---|---|---|---|---|---|---|---|---|---|---|---|
| | | | Mean Optical Densities | | | | | Percent Growth | | | | | | | |
| Panel/Cell Line | Zero | Ctrl | -8.3 | -7.3 | -6.3 | -5.3 | -4.3 | -8.3 | -7.3 | -6.3 | -5.3 | -4.3 | GI50 | TGI | LC50 |
| **Leukemia** | | | | | | | | | | | | | | | |
| CCRF-CEM | 0.499 | 2.474 | 2.658 | 2.854 | 2.352 | 0.573 | 0.363 | 109 | 119 | 94 | 4 | -27 | 1.53E-6 | 6.59E-6 | > 5.00E-5 |
| HL-60(TB) | 0.561 | 2.401 | 2.393 | 2.538 | 2.590 | 0.468 | 0.319 | 100 | 107 | 110 | -17 | -43 | 1.49E-6 | 3.70E-6 | > 5.00E-5 |
| K-562 | 0.184 | 2.006 | 1.966 | 2.071 | 1.772 | 0.229 | 0.230 | 98 | 104 | 87 | 2 | 2 | 1.37E-6 | > 5.00E-5 | > 5.00E-5 |
| MOLT-4 | 0.573 | 2.847 | 2.867 | 3.032 | 2.846 | 0.765 | 0.289 | 101 | 108 | 100 | 8 | -50 | 1.76E-6 | 6.99E-6 | > 5.00E-5 |
| RPMI-8226 | 0.997 | 3.005 | 2.963 | 2.935 | 2.819 | 0.701 | 0.572 | 98 | 96 | 91 | -30 | -43 | 1.09E-6 | 2.83E-6 | > 5.00E-5 |
| SR | 0.550 | 2.546 | 2.265 | 2.361 | 2.255 | 0.865 | 0.385 | 86 | 91 | 85 | 16 | -30 | 1.61E-6 | 1.10E-5 | > 5.00E-5 |
| **Non-Small Cell Lung Cancer** | | | | | | | | | | | | | | | |
| A549/ATCC | 0.872 | 2.984 | 2.828 | 2.880 | 2.830 | 1.339 | 0.966 | 93 | 95 | 93 | 22 | 4 | 2.01E-6 | > 5.00E-5 | > 5.00E-5 |
| EKVX | 0.803 | 2.244 | 2.172 | 2.243 | 2.207 | 1.087 | 0.886 | 95 | 100 | 97 | 20 | 6 | 2.04E-6 | > 5.00E-5 | > 5.00E-5 |
| HOP-62 | 0.902 | 2.681 | 2.633 | 2.696 | 2.671 | 0.977 | 0.473 | 97 | 101 | 99 | 4 | -48 | 1.65E-6 | 6.03E-6 | > 5.00E-5 |
| HOP-92 | 1.688 | 2.213 | 2.100 | 2.191 | 2.112 | 1.171 | 1.130 | 79 | 96 | 81 | -31 | -33 | 9.45E-7 | 2.65E-6 | > 5.00E-5 |
| NCI-H226 | 1.682 | 2.750 | 2.692 | 2.723 | 2.633 | 1.211 | 0.317 | 95 | 97 | 89 | -28 | -81 | 1.08E-6 | 2.88E-6 | 1.30E-5 |
| NCI-H322M | 0.792 | 2.257 | 2.164 | 2.153 | 2.100 | 0.857 | 0.870 | 94 | 93 | 89 | 4 | 5 | 1.45E-6 | > 5.00E-5 | > 5.00E-5 |
| NCI-H460 | 0.230 | 2.330 | 2.395 | 2.419 | 2.374 | 0.363 | 0.192 | 103 | 104 | 102 | 6 | -17 | 1.75E-6 | 9.46E-6 | > 5.00E-5 |
| NCI-H522 | 1.048 | 2.901 | 2.741 | 2.753 | 2.649 | 0.773 | 0.685 | 91 | 92 | 86 | -26 | -35 | 1.05E-6 | 2.92E-6 | > 5.00E-5 |
| **Colon Cancer** | | | | | | | | | | | | | | | |
| COLO 205 | 0.480 | 2.117 | 2.111 | 2.165 | 1.995 | 0.657 | 0.105 | 100 | 103 | 93 | 11 | -78 | 1.66E-6 | 6.62E-6 | 2.41E-5 |
| HCC-2998 | 0.737 | 2.487 | 2.328 | 2.532 | 2.447 | 0.918 | 0.693 | 91 | 103 | 98 | 10 | -6 | 1.76E-6 | 2.14E-5 | > 5.00E-5 |
| HCT-116 | 0.228 | 2.316 | 2.348 | 2.239 | 2.168 | 0.264 | 0.046 | 102 | 96 | 93 | 2 | -80 | 1.48E-6 | 5.25E-6 | 2.15E-5 |
| HCT-15 | 0.382 | 2.537 | 2.438 | 2.454 | 2.311 | 0.475 | 0.268 | 95 | 96 | 89 | 4 | -30 | 1.45E-6 | 6.68E-6 | > 5.00E-5 |
| HT29 | 0.369 | 2.401 | 2.300 | 2.420 | 2.344 | 0.697 | 0.204 | 95 | 101 | 97 | 16 | -45 | 1.91E-6 | 9.20E-6 | > 5.00E-5 |
| KM12 | 0.696 | 3.217 | 3.182 | 3.189 | 3.138 | 0.969 | 0.578 | 99 | 99 | 97 | 11 | -17 | 1.75E-6 | 1.23E-5 | > 5.00E-5 |
| SW-620 | 0.267 | 1.981 | 2.000 | 2.072 | 1.863 | 0.467 | 0.272 | 101 | 105 | 93 | 12 | 0 | 1.69E-6 | > 5.00E-5 | > 5.00E-5 |
| **CNS Cancer** | | | | | | | | | | | | | | | |
| SF-268 | 0.898 | 2.630 | 2.502 | 2.595 | 2.531 | 1.088 | 1.000 | 93 | 98 | 94 | 11 | 6 | 1.70E-6 | > 5.00E-5 | > 5.00E-5 |
| SF-295 | 0.951 | 2.656 | 2.618 | 2.642 | 2.633 | 0.575 | 0.371 | 98 | 99 | 99 | -40 | -61 | 1.12E-6 | 2.59E-6 | 1.53E-5 |
| SF-539 | 0.706 | 2.408 | 2.330 | 2.348 | 2.364 | 1.059 | 0.669 | 95 | 96 | 97 | 21 | -5 | 2.08E-6 | 3.14E-5 | > 5.00E-5 |
| SNB-19 | 0.553 | 1.944 | 1.861 | 1.901 | 1.850 | 0.863 | 0.774 | 94 | 97 | 93 | 22 | 16 | 2.03E-6 | > 5.00E-5 | > 5.00E-5 |
| SNB-75 | 1.689 | 2.686 | 2.464 | 2.475 | 2.445 | 1.646 | 1.517 | 78 | 79 | 76 | -3 | -10 | 1.07E-6 | 4.64E-6 | > 5.00E-5 |
| U251 | 0.706 | 2.843 | 2.747 | 2.778 | 2.498 | 1.004 | 0.639 | 96 | 97 | 84 | 14 | -9 | 1.52E-6 | 1.97E-5 | > 5.00E-5 |
| **Melanoma** | | | | | | | | | | | | | | | |
| LOX IMVI | 0.320 | 1.903 | 1.901 | 1.894 | 1.856 | 1.219 | 0.137 | 100 | 99 | 97 | 57 | -57 | 5.74E-6 | 1.57E-5 | 4.31E-5 |
| MALME-3M | 0.621 | 1.226 | 1.222 | 1.233 | 1.232 | 0.820 | 0.445 | 99 | 101 | 101 | 33 | -28 | 2.80E-6 | 1.72E-5 | > 5.00E-5 |
| M14 | 0.560 | 2.343 | 2.264 | 2.316 | 2.198 | 0.963 | 0.468 | 96 | 98 | 92 | 23 | -17 | 2.01E-6 | 1.89E-5 | > 5.00E-5 |
| MDA-MB-435 | 0.499 | 2.308 | 2.241 | 2.246 | 2.188 | 1.167 | 0.366 | 96 | 97 | 93 | 37 | -27 | 2.93E-6 | 1.90E-5 | > 5.00E-5 |
| SK-MEL-2 | 1.722 | 2.753 | 2.667 | 2.760 | 2.724 | 0.911 | 0.470 | 92 | 101 | 97 | -47 | -73 | 1.06E-6 | 2.36E-6 | 6.49E-6 |
| SK-MEL-28 | 0.625 | 1.992 | 2.016 | 2.036 | 1.909 | 0.895 | 0.536 | 102 | 103 | 94 | 20 | -14 | 1.95E-6 | 1.91E-5 | > 5.00E-5 |
| SK-MEL-5 | 0.912 | 3.297 | 3.309 | 3.296 | 3.245 | 0.434 | 0.311 | 101 | 100 | 98 | -52 | -66 | 1.04E-6 | 2.24E-6 | 4.81E-6 |
| UACC-257 | 1.294 | 2.910 | 2.762 | 2.827 | 2.704 | 1.455 | 0.914 | 91 | 95 | 87 | 10 | -29 | 1.52E-6 | 8.95E-6 | > 5.00E-5 |
| UACC-62 | 0.837 | 2.868 | 2.748 | 2.818 | 2.794 | 0.776 | 0.454 | 94 | 98 | 96 | -7 | -46 | 1.40E-6 | 4.25E-6 | > 5.00E-5 |
| **Ovarian Cancer** | | | | | | | | | | | | | | | |
| IGROV1 | 0.520 | 2.317 | 2.232 | 2.316 | 2.240 | 1.097 | 0.643 | 95 | 100 | 96 | 32 | 7 | 2.62E-6 | > 5.00E-5 | > 5.00E-5 |
| OVCAR-3 | 0.601 | 2.068 | 2.110 | 2.112 | 1.964 | 0.895 | 0.660 | 103 | 103 | 93 | 20 | 4 | 1.94E-6 | > 5.00E-5 | > 5.00E-5 |
| OVCAR-4 | 0.656 | 1.752 | 1.696 | 1.758 | 1.662 | 0.851 | 0.794 | 95 | 101 | 92 | 18 | 13 | 1.83E-6 | > 5.00E-5 | > 5.00E-5 |
| OVCAR-5 | 0.703 | 1.878 | 1.802 | 1.831 | 1.796 | 1.282 | 0.915 | 93 | 96 | 93 | 49 | 18 | 4.81E-6 | > 5.00E-5 | > 5.00E-5 |
| OVCAR-8 | 0.308 | 1.588 | 1.591 | 1.598 | 1.523 | 0.407 | 0.288 | 100 | 101 | 95 | 8 | -7 | 1.64E-6 | 1.72E-5 | > 5.00E-5 |
| NCI/ADR-RES | 0.581 | 1.996 | 1.976 | 2.070 | 1.957 | 0.555 | 0.497 | 99 | 105 | 97 | -5 | -14 | 1.45E-6 | 4.51E-6 | > 5.00E-5 |
| SK-OV-3 | 1.170 | 2.256 | 2.230 | 2.288 | 2.207 | 1.068 | 0.384 | 98 | 103 | 95 | -9 | -67 | 1.36E-6 | 4.12E-6 | 2.54E-5 |
| **Renal Cancer** | | | | | | | | | | | | | | | |
| 786-0 | 0.478 | 2.335 | 2.178 | 2.300 | 2.221 | 0.835 | 0.411 | 92 | 98 | 94 | 19 | -14 | 1.93E-6 | 1.89E-5 | > 5.00E-5 |
| A498 | 1.867 | 2.520 | 2.543 | 2.608 | 2.566 | 1.712 | 1.665 | 103 | 113 | 107 | -8 | -11 | 1.56E-6 | 4.23E-6 | > 5.00E-5 |
| ACHN | 0.260 | 1.404 | 1.423 | 1.455 | 1.404 | 0.698 | 0.312 | 102 | 104 | 100 | 38 | 5 | 3.23E-6 | > 5.00E-5 | > 5.00E-5 |
| CAKI-1 | 0.540 | 1.940 | 1.799 | 1.820 | 1.710 | 0.956 | 0.459 | 90 | 91 | 84 | 30 | -15 | 2.10E-6 | 2.30E-5 | > 5.00E-5 |
| RXF 393 | 0.825 | 1.459 | 1.450 | 1.524 | 1.444 | 0.706 | 0.380 | 99 | 110 | 98 | -14 | -54 | 1.33E-6 | 3.72E-6 | 3.96E-5 |
| SN12C | 0.572 | 2.204 | 2.124 | 2.204 | 2.100 | 0.590 | 0.522 | 95 | 100 | 94 | 1 | -9 | 1.48E-6 | 6.43E-6 | > 5.00E-5 |
| TK-10 | 1.383 | 2.246 | 2.101 | 2.162 | 2.256 | 1.299 | 1.246 | 83 | 90 | 101 | -6 | -10 | 1.50E-6 | 4.39E-6 | > 5.00E-5 |
| UO-31 | 0.610 | 2.003 | 1.779 | 1.839 | 1.766 | 1.362 | 0.292 | 84 | 88 | 83 | 54 | -52 | 5.45E-6 | 1.61E-5 | 4.77E-5 |
| **Prostate Cancer** | | | | | | | | | | | | | | | |
| PC-3 | 0.625 | 2.607 | 2.575 | 2.672 | 2.498 | 0.594 | 0.558 | 98 | 103 | 94 | -5 | -11 | 1.40E-6 | 4.45E-6 | > 5.00E-5 |
| DU-145 | 0.442 | 2.024 | 2.095 | 2.146 | 2.062 | 0.946 | 0.641 | 104 | 108 | 102 | 32 | 13 | 2.76E-6 | > 5.00E-5 | > 5.00E-5 |
| **Breast Cancer** | | | | | | | | | | | | | | | |
| MCF7 | 0.461 | 2.309 | 2.134 | 2.205 | 2.201 | 0.502 | 0.445 | 90 | 94 | 94 | 2 | -3 | 1.51E-6 | 1.23E-5 | > 5.00E-5 |
| MDA-MB-231/ATCC | 0.540 | 1.468 | 1.476 | 1.520 | 1.457 | 0.593 | 0.439 | 101 | 106 | 99 | 6 | -19 | 1.67E-6 | 8.52E-6 | > 5.00E-5 |
| HS 578T | 1.734 | 2.745 | 2.665 | 2.720 | 2.685 | 1.454 | 1.521 | 92 | 97 | 94 | -16 | -12 | 1.25E-6 | 3.57E-6 | > 5.00E-5 |
| BT-549 | 1.658 | 2.960 | 2.940 | 2.948 | 2.974 | 1.354 | 0.985 | 98 | 99 | 101 | -18 | -41 | 1.34E-6 | 3.51E-6 | > 5.00E-5 |
| T-47D | 0.966 | 2.279 | 2.161 | 2.221 | 2.193 | 1.069 | 0.876 | 91 | 96 | 93 | 8 | -9 | 1.61E-6 | 1.43E-5 | > 5.00E-5 |
| MDA-MB-468 | 0.790 | 1.327 | 1.317 | 1.347 | 1.252 | 0.736 | 0.613 | 98 | 104 | 86 | -7 | -22 | 1.22E-6 | 4.21E-6 | > 5.00E-5 |

**Fig. S2:** Cytotoxicity data derived from the NCI-60 panel of the National Cancer Institute (NCI) Developmental Therapeutics Program (five-dose-SRB-assay, 5 nM to 50 µM). **(A)** Effect of AETX on cell growth in % on various cancer cell lines. **(B)** Individual data for every cancer cell line. Results are summarized according to cell line origin.GI50: growth inhibition 50%, TGI: total growth inhibition, LC50: lethal concentration 50%.

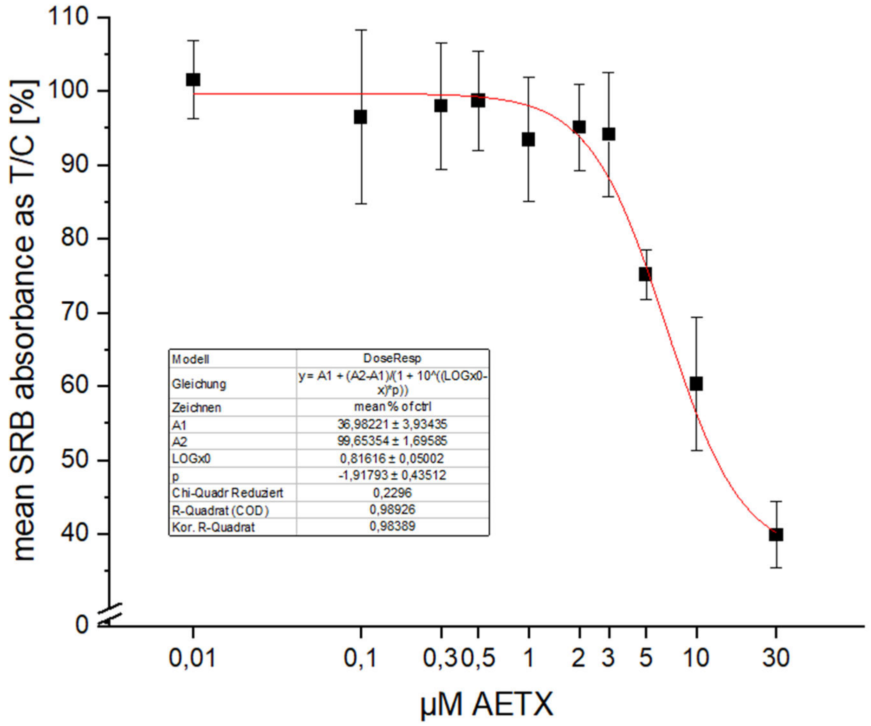

**Fig. S3:** Mean SRB absorbance of AETX-treated HeLa cells normalized to the control. Whiskers indicate standard deviation. Red line: calculated fit. $EC_{50}$ 6.5 µM.

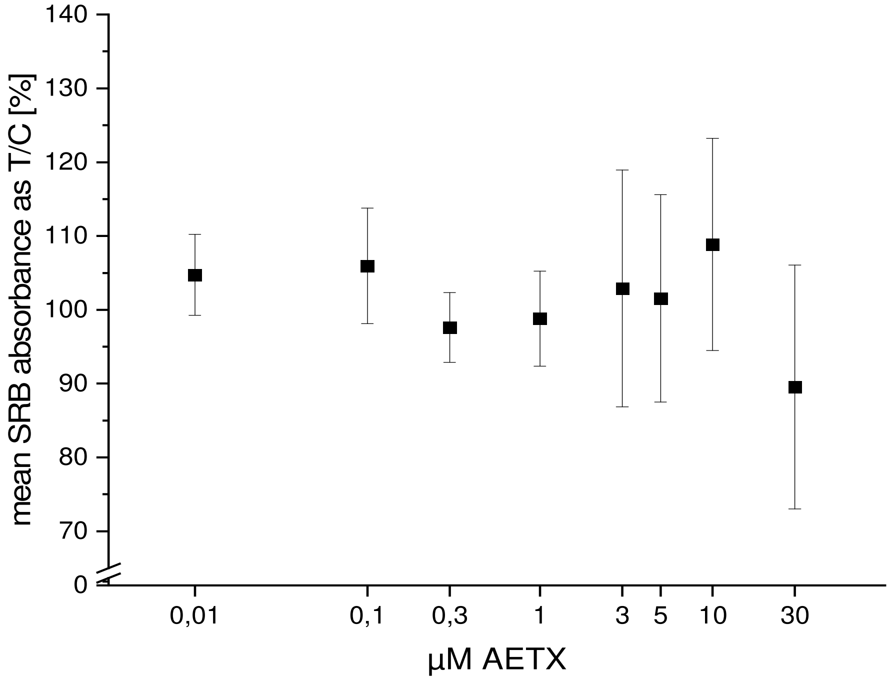

**Fig. S4:** Mean SRB absorbance of AETX-treated fibroblasts normalized to the control. Whiskers indicate standard deviation. A fit could not be calculated. $EC_{50}$ > 30 µM.

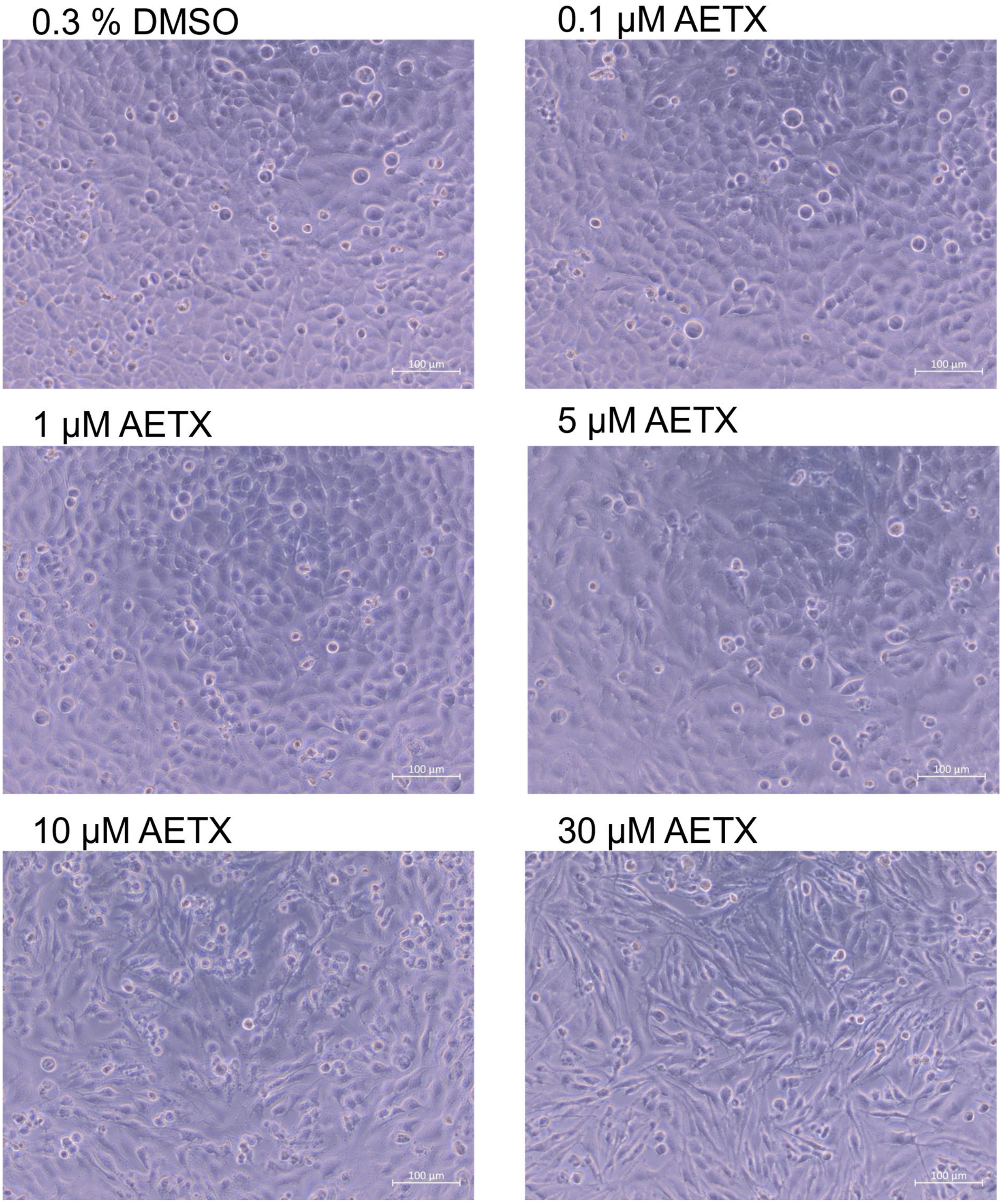

**Fig. S5:** Phase contrast microscopy images of morphological changes of HeLa cells incubated with 0.3% DMSO, 0.1, 1, 5, 10, and 30 µM AETX for 24 h. Scale bar 100 µm.

A

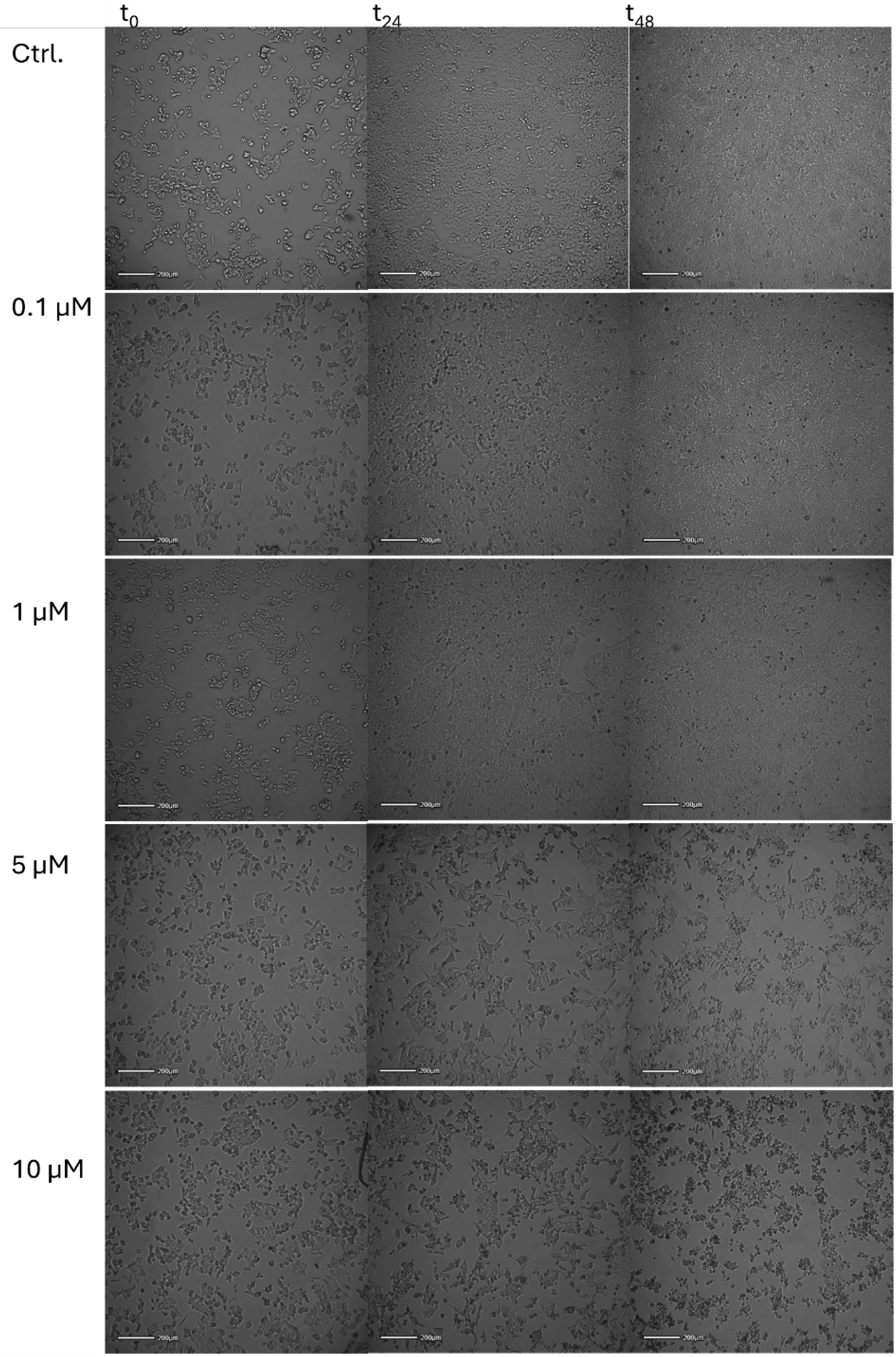

**B**

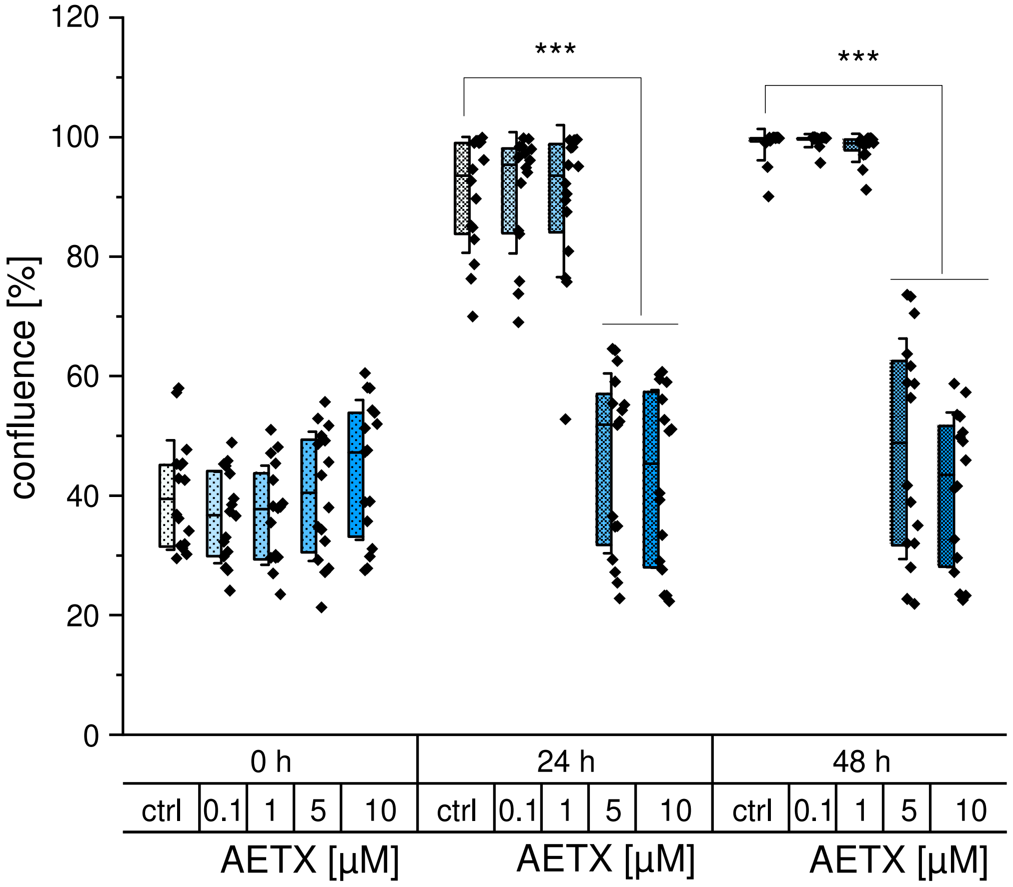

**Fig. S6:** Effect of AETX on HCT116 cell confluency. **(A)** Brightfield microscopy images showing confluency of AETX-treated HCT116 cells directly (t0), 24 h (t24) and 48 h (t48) after treatment. Scale bar 200 µm. **(B)** Confluency of AETX-treated HCT116 cells at different timepoints after treatment in % of covered area acquired from brightfield microscopy images. Cells were treated with 0.1% DMSO (ctrl), 0.1, 1, 5, or 10 µM AETX. Statistically significant difference to the control indicated with *** $p < 0.001$, obtained with Student's t-test. Data based on four independent biological replicates.

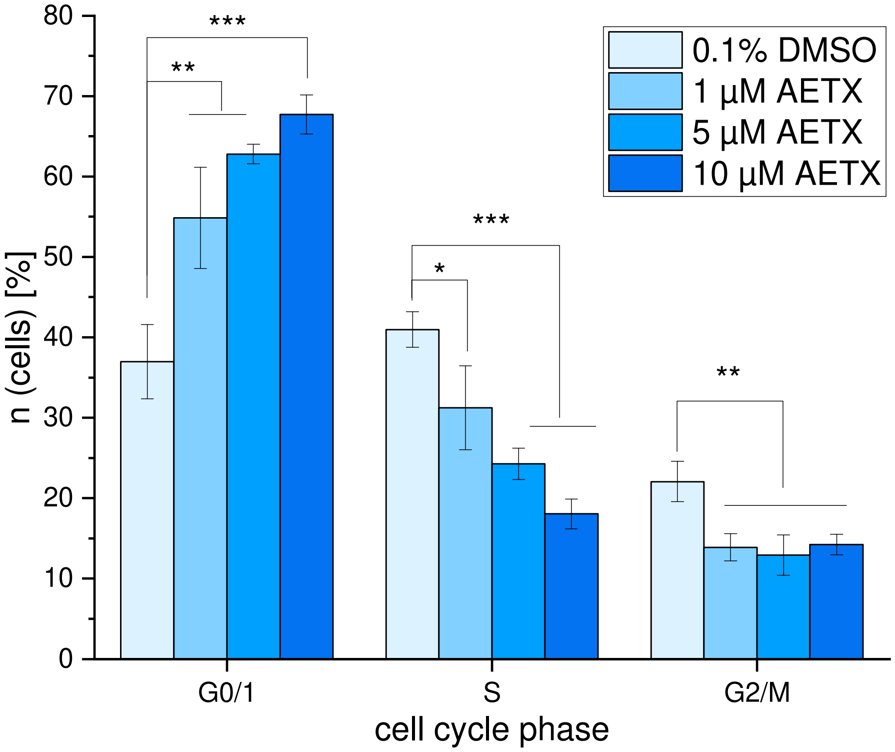

**Fig. S7:** Distribution of cell cycle phases in HCT116 cell populations treated with 0.1% DMSO, 1, 5, or 10 µM AETX for 24 h. Statistically significant difference to the control indicated with *p<0.05, **p<0.01, ***p<0.001, obtained with Student's t-test. Data results from four independent biological replicates.

**A**

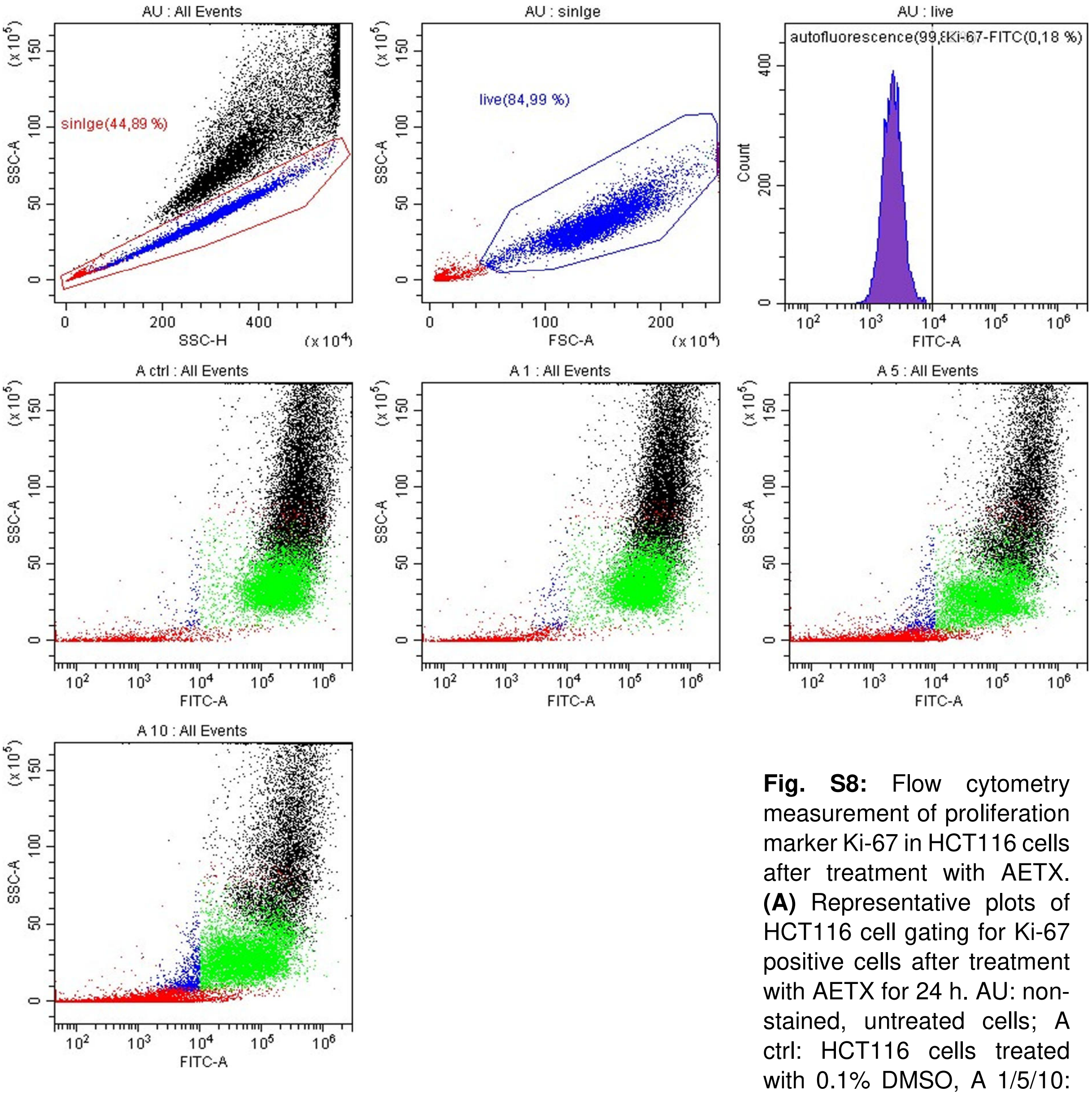

**B**

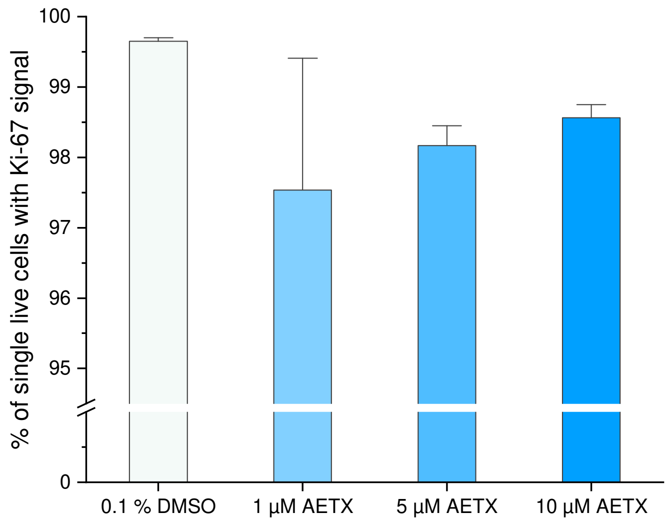

**Fig. S8:** Flow cytometry measurement of proliferation marker Ki-67 in HCT116 cells after treatment with AETX. **(A)** Representative plots of HCT116 cell gating for Ki-67 positive cells after treatment with AETX for 24 h. AU: non-stained, untreated cells; A ctrl: HCT116 cells treated with 0.1% DMSO, A 1/5/10: HCT116 cells treated with 1, 5, or 10 µM AETX. Gates: red: cell singlets, blue: healthy cells, green: Ki-67-positive cells, black: cells which are excluded from the gates. **(B)** HCT116 cells with Ki-67 signal after 24 h of treatment with 0.1% DMSO, 1, 5, or 10 µM AETX. No statistically significant difference to the control, obtained with Mann-Whitney test. Data obtained from four independent biological replicates. Whiskers indicate standard deviation (1x).

**Metabolomics**

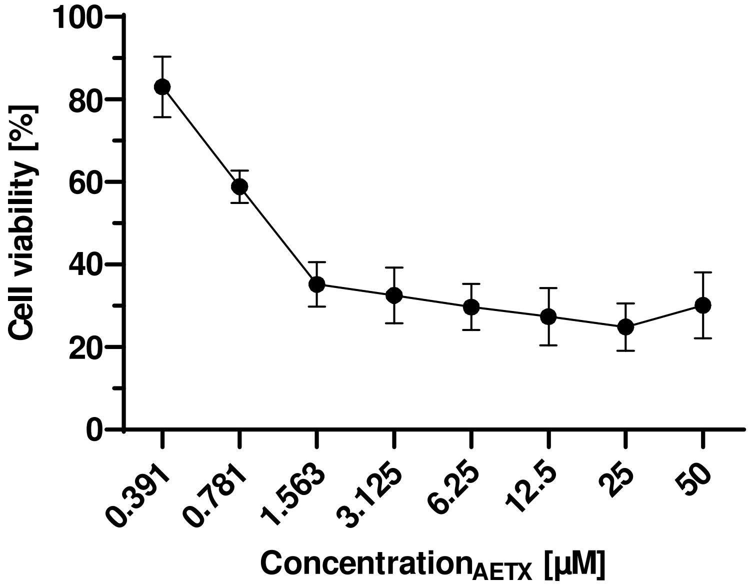

**Fig. S9:** Mean viability of PC-3 cells treated with several concentrations of AETX as compared to the control. Data determined under use of the MTT assay. Whiskers show standard deviation (1x).

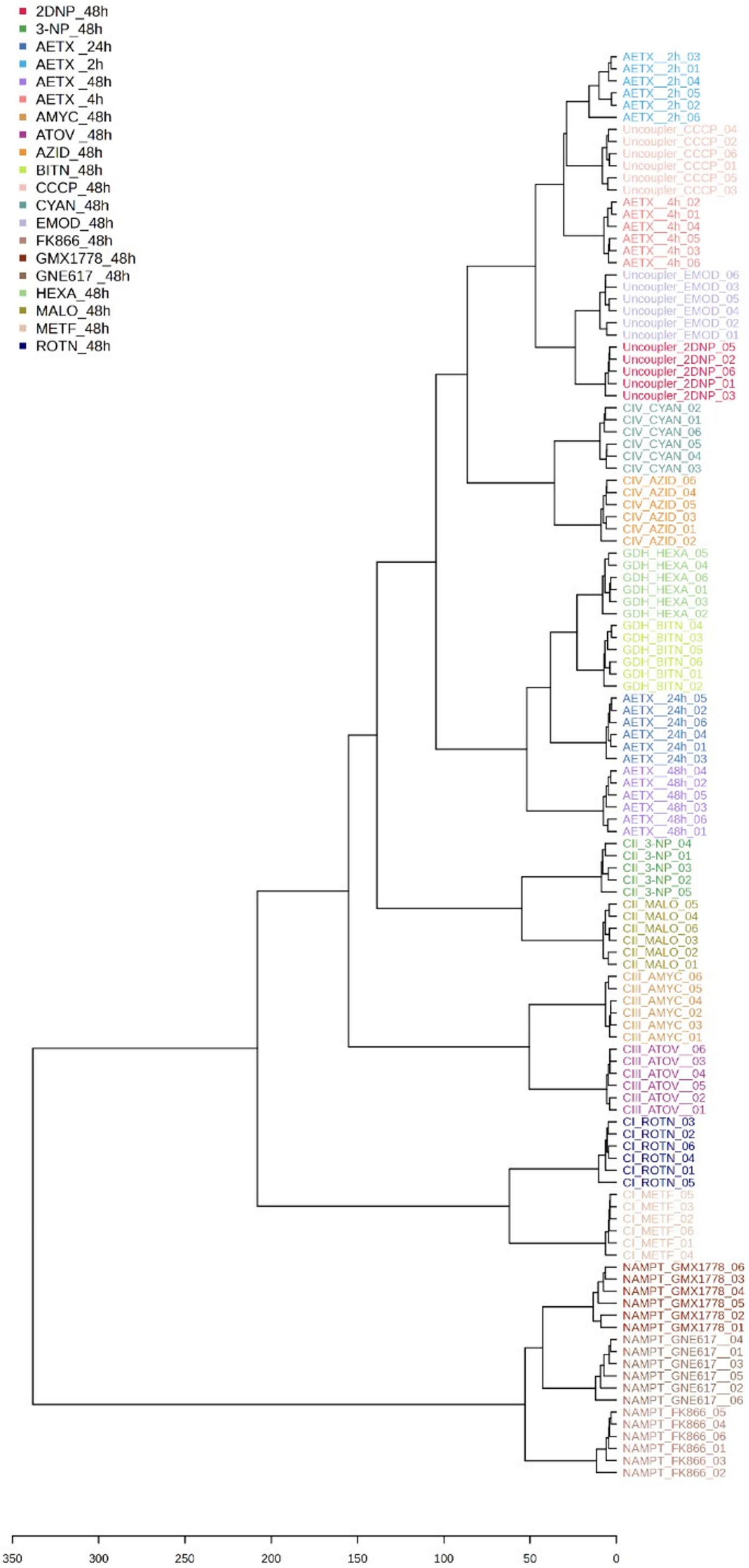

**Fig. S10:** Hierarchical cluster analysis of metabolic profiles induced by OXPHOS inhibitors, NAMPT inhibitors, GDH inhibitors and AETX based on the modulation of the metabolism of prostate cancer cells (PC-3). Data aggregated from six replicates for each experiment.
<u>Compounds:</u>
2DNP - 2,4-dinitrophenol
3-NP - 3-nitropropionic acid
AMYC – antimycin A
ATOV – atovaquone
AZID - sodium azide
BITN – bithionol
CCCP - carbonyl cyanide chlorophenylhydrazone
CYAN - potassium cyanide
EMOD – emodin
FK866 - FK866
GMX - GMX1778
GNE - GNE-617
HEXA – hexachlorophene
MALO - malonic acid
METF – metformin
ROTN - rotenone.
<u>MoA:</u>
CPLX I - complex I
CPLX II - complex II
CPLX III - complex III
CPLX IV - complex IV
GDH - glutamate dehydrogenase
NAMPT - nicotinamide phosphorribosyltransferase
Uncoupler - uncoupling of oxidative phosphorylation.

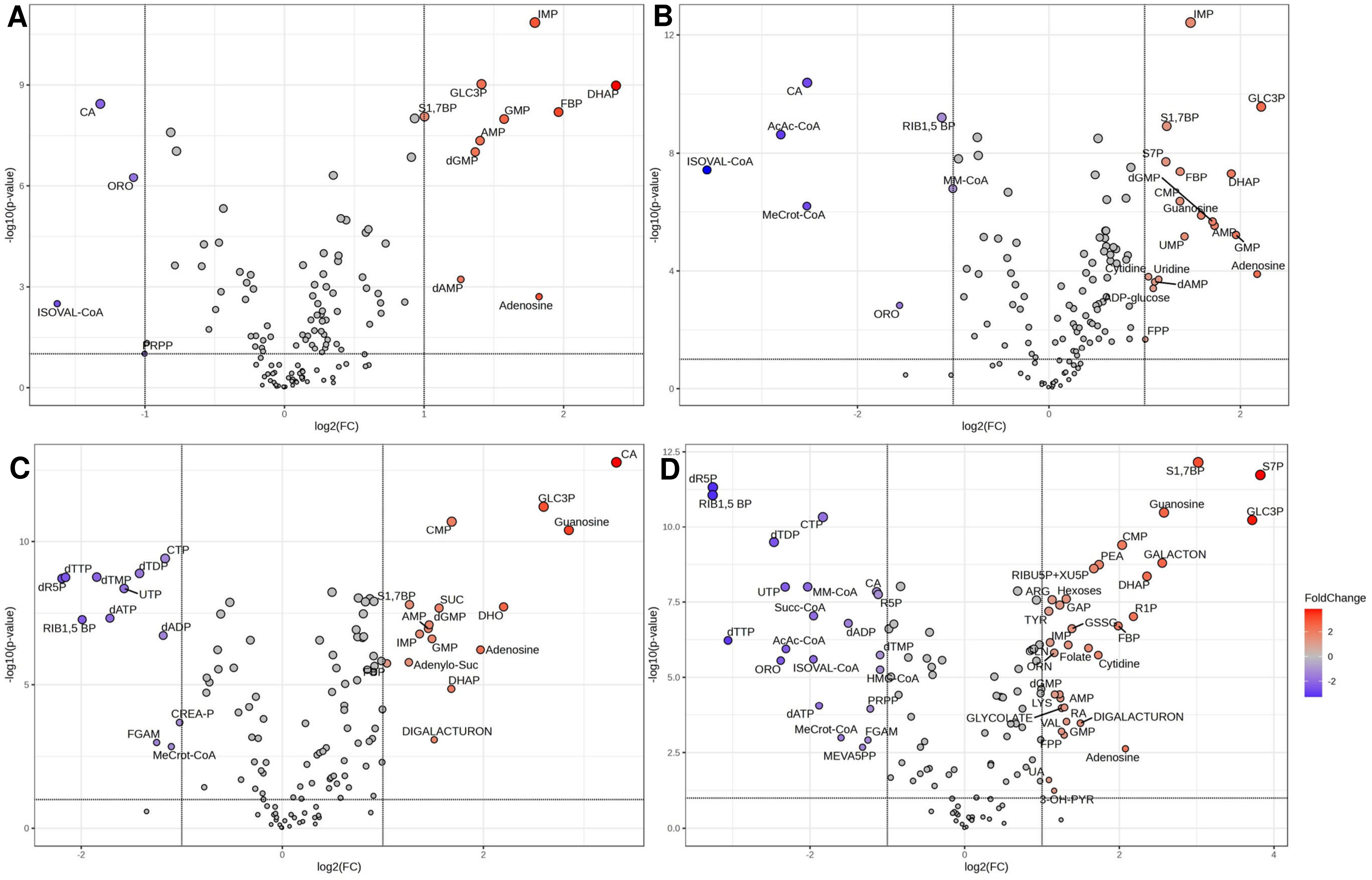

**Fig. S11:** Log fold changes between metabolites of PC3 cells incubated for 2 h **(A)**, 4 h **(B)**, 24 h **(C)**, and 48 h **(D)** with 0.74 µM AETX or solvent control. Statistical significance indicated in red (higher than control) or blue (lower than control). For abbreviations see supplementary metabolomics data (Excel file).

### Chemistry

### pKa and logP

Yasuda-Shedlovsky Result

Multiset name: **UV-metric psKa AETX** Instrument ID: **T310018**
Analyst: **RR**
Quality: **Good**
Filename: **E:\AETX\Experimental\AETX_UV-metric psKa_MeOH - Multiset.t3r**

#### Yasuda-Shedlovsky result

| Extrapolation type | pKa0% | SD | Intercept | Slope | $R^2$ | Ionic strength | Temperature |
|---|---|---|---|---|---|---|---|
| Yasuda-Shedlovsky | 6.87 | ±0.03 | 9.70 | -85.0455 | 0.9939 | 0.163 M | 24.9°C |

#### Component assay results

| Titration | Methanol weight% | Direction | Result type | Dielectric constant | [H2O] | Ionic strength | Temperature | | psKa 1 |
|---|---|---|---|---|---|---|---|---|---|
| 23I-12006 Points 48 to 91 | 52.65 % | Up | UV-metric pKa | 55.1 | 23.0 M | 0.169 M | 25.0°C | ☑ | 6.79 |
| 23I-12013 Points 46 to 88 | 56.46 % | Up | UV-metric pKa | 53.3 | 20.9 M | 0.169 M | 25.0°C | ☑ | 6.79 |
| 23I-13004 Points 4 to 44 | 58.64 % | Up | UV-metric pKa | 52.3 | 19.8 M | 0.157 M | 24.9°C | ☑ | 6.77 |
| 23I-12006 Points 4 to 46 | 62.91 % | Up | UV-metric pKa | 50.2 | 17.6 M | 0.157 M | 24.9°C | ☑ | 6.76 |

#### Graphs

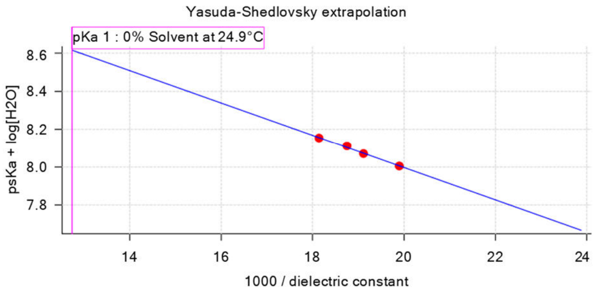

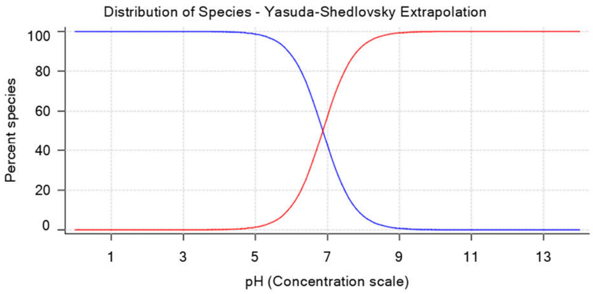

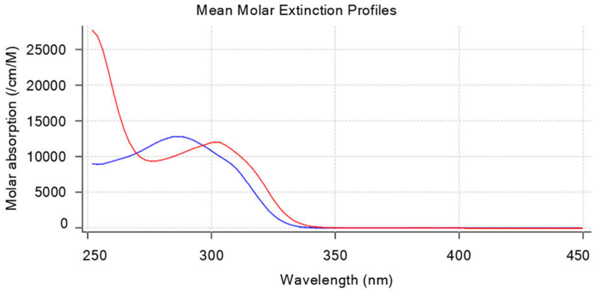

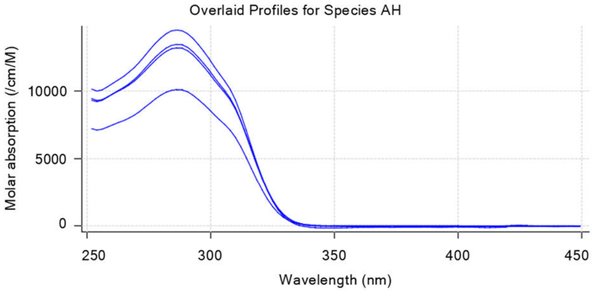

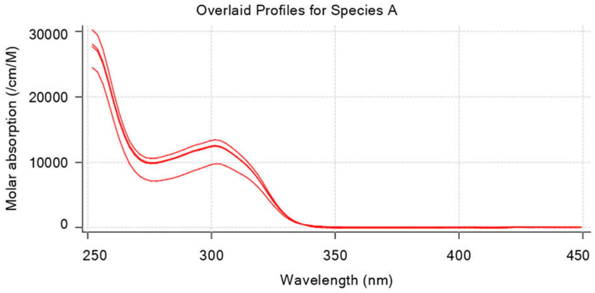

### UV-metric psKa 23I-12006 Assay 1 of 3 Quality: Good

#### Calibration Settings

**Setting Value Date/Time changed Imported from**
Four-Plus alpha 0.166 12/09/2023 12:34:55 H:\Data 2023\T310018\September\23I-12003_Blank standardisation.t3r
Four-Plus S 0.9994 12/09/2023 12:34:55 H:\Data 2023\T310018\September\23I-12003_Blank standardisation.t3r Four-Plus jH 0.6 12/09/2023 12:34:55 H:\Data 2023\T310018\September\23I-12003_Blank standardisation.t3r

Multiset name: **UV-metric psKa AETX** Instrument ID: **T310018**
Analyst: **RR**
Quality: **Good**
Filename: **E:\AETX\Experimental\AETX_UV-metric psKa_MeOH - Multiset.t3r**

### Calibration Settings (continued)

| Setting | Value | Date/Time changed | Imported from |
|---|---|---|---|
| Four-Plus jOH | -0.6 | 12/09/2023 12:34:55 | H:\Data 2023\T310018\September\23I-12003_Blank standardisation.t3r |
| Base concentration factor | 1.025 | 12/09/2023 12:34:55 | H:\Data 2023\T310018\September\KHP Multiset 1st September 2023_PASS.t3r |
| Acid concentration factor | 0.988 | 12/09/2023 12:34:55 | H:\Data 2023\T310018\September\23I-12003_Blank standardisation.t3r |

### UV-metric psKa_Low 23I-12013 Assay 2 of 3 Quality: Good

#### Calibration Settings

| Setting | Value | Date/Time changed | Imported from |
|---|---|---|---|
| Four-Plus alpha | 0.166 | 12/09/2023 17:14:08 | H:\Data 2023\T310018\September\23I-12003_Blank standardisation.t3r |
| Four-Plus S | 0.9994 | 12/09/2023 17:14:08 | H:\Data 2023\T310018\September\23I-12003_Blank standardisation.t3r |
| Four-Plus jH | 0.6 | 12/09/2023 17:14:08 | H:\Data 2023\T310018\September\23I-12003_Blank standardisation.t3r |
| Four-Plus jOH | -0.6 | 12/09/2023 17:14:08 | H:\Data 2023\T310018\September\23I-12003_Blank standardisation.t3r |
| Base concentration factor | 1.025 | 12/09/2023 17:14:08 | H:\Data 2023\T310018\September\KHP Multiset 1st September 2023_PASS.t3r |
| Acid concentration factor | 0.988 | 12/09/2023 17:14:08 | H:\Data 2023\T310018\September\23I-12003_Blank standardisation.t3r |

### UV-metric psKa_Low 60MeOH 23I-13004 Assay 3 of 3 Quality: Good

#### Calibration Settings

| Setting | Value | Date/Time changed | Imported from |
|---|---|---|---|
| Four-Plus alpha | 0.143 | 13/09/2023 12:48:12 | H:\Data 2023\T310018\September\23I-13003_Blank standardisation.t3r |
| Four-Plus S | 0.9997 | 13/09/2023 12:48:12 | H:\Data 2023\T310018\September\23I-13003_Blank standardisation.t3r |
| Four-Plus jH | 0.4 | 13/09/2023 12:48:12 | H:\Data 2023\T310018\September\23I-13003_Blank standardisation.t3r |
| Four-Plus jOH | -0.4 | 13/09/2023 12:48:12 | H:\Data 2023\T310018\September\23I-13003_Blank standardisation.t3r |
| Base concentration factor | 1.025 | 13/09/2023 12:48:12 | H:\Data 2023\T310018\September\KHP Multiset 1st September 2023_PASS.t3r |
| Acid concentration factor | 0.987 | 13/09/2023 12:48:12 | H:\Data 2023\T310018\September\23I-13003_Blank standardisation.t3r |

**Fig. S12:** Report of the UV-metric p$K_a$ assessment for AETX.

pH-metric Result

| | | | |
|---|---|---|---|
| Sample name: | **AETX** | Experiment start time: | **29/09/2023 18:10:55** |
| Assay name: | **pH-metric high logP** | Analyst: | **RR** |
| Assay ID: | **23I-29010** | Instrument ID: | **T311054** |
| Quality: | **Good** | | |
| Filename: | **E:\AETX\Experimental\23I-29010_AETX_pH-metric high logP.t3r** | | |

### pH-metric Result

logP (neutral XH) 4.74 ±0.07 (n=50)

### Sample logD values

| pH | AETX logD | Comment |
|---|---|---|
| 1.000 | 4.74 | |
| 1.200 | 4.74 | Stomach pH |
| 2.000 | 4.74 | |
| 3.000 | 4.74 | |
| 4.000 | 4.74 | |
| 5.000 | 4.74 | |
| 6.000 | 4.69 | |
| 6.500 | 4.59 | |
| 7.000 | 4.37 | |
| 7.400 | 4.10 | Blood pH |
| 8.000 | 3.58 | |
| 9.000 | 2.61 | |
| 10.000 | 1.61 | |
| 11.000 | 0.61 | |
| 12.000 | -0.39 | |

### Graphs

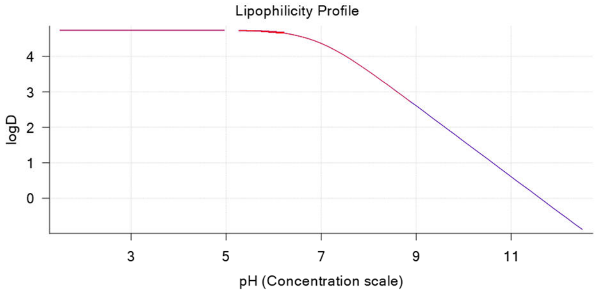

### Calibration Settings

**Setting** **Value Date/Time changed Imported from**
Four-Plus alpha 0.180 29/09/2023 18:10:55 H:\Data 2023\T311054\September\23I-29004_Blank standardisation.t3r
Four-Plus S 0.9930 29/09/2023 18:10:55 H:\Data 2023\T311054\September\23I-29004_Blank standardisation.t3r
Four-Plus jH 0.8 29/09/2023 18:10:55 H:\Data 2023\T311054\September\23I-29004_Blank standardisation.t3r Four-
Plus jOH 0.0 29/09/2023 18:10:55 H:\Data 2023\T311054\September\23I-29004_Blank standardisation.t3r Base
concentration factor 1.020 29/09/2023 18:10:55 H:\Data 2023\T311054\September\KHP Multiset 1st September
2023_PASS.t3r
Acid concentration factor 0.993 29/09/2023 18:10:55 H:\Data 2023\T311054\September\23I-29004_Blank standardisation.t3r

Reported at: 01/11/2023 13:58:35 Page 1 of 1

**Fig. S13:** Report of the pH-metric logP assessment for AETX.

**Cell biology**

**Seahorse experiments**

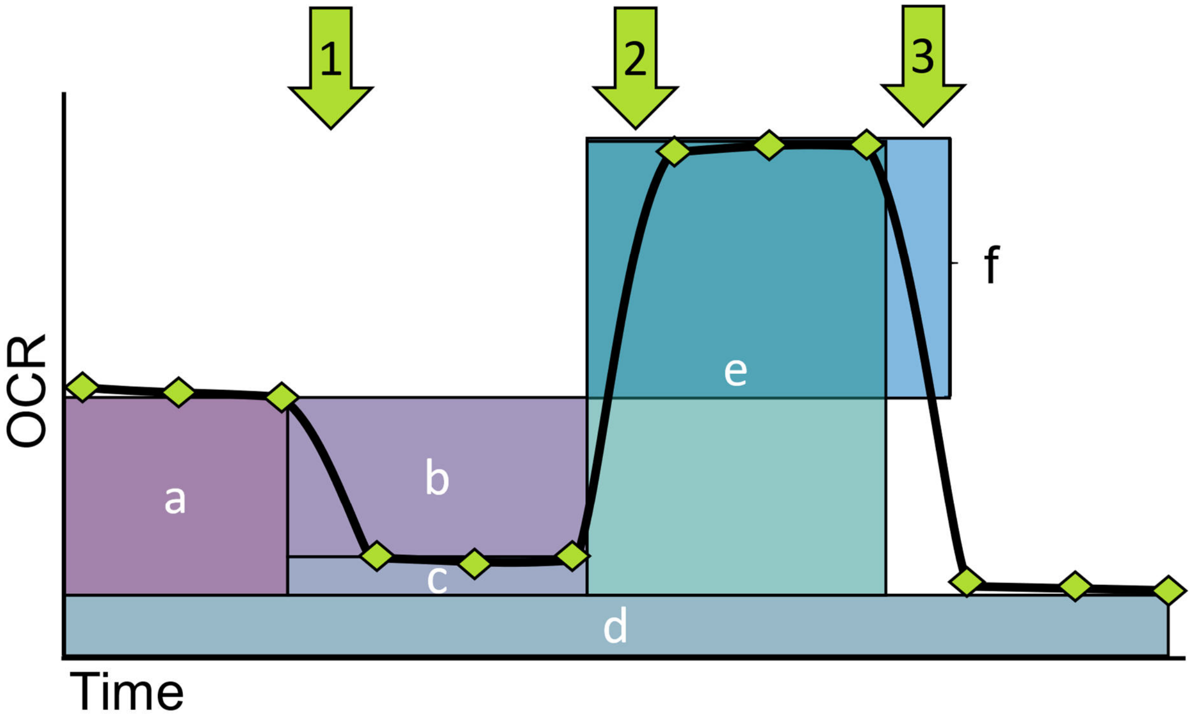

**Fig. S14:** Sketch of a model oxygen consumption rate assay with the Seahorse FX analyzer. OCR: Oxygen Consumption Rate. Green diamonds indicate single measurements. Green arrows indicate the addition of compounds in the following order: 1: oligomycin to inhibit FoF1-ATPase, 2: FCCP to stimulate respiration, 3: rotenone/antimycin A + Hoechst 33342 to inhibit respiratory chain activity and to stain DNA for subsequent normalization on cell count. Areas indicate endpoints accessible through this approach: a: basal respiration, b: mitochondrial ATP production, c: proton leak, d: non-mitochondrial oxygen consumption, e: maximal respiration, f: spare respiratory capacity.

**PER**

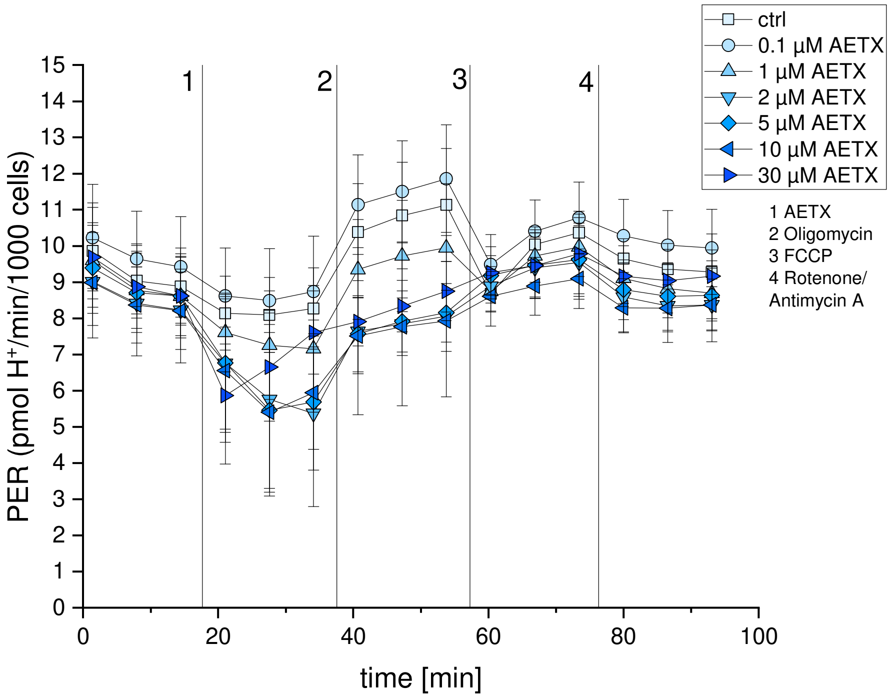

**Fig. S15:** Proton Efflux Rate (PER) measurement of fibroblasts after acute stimulation with AETX. □: solvent control (0.1% DMSO), ○: 0.1 µM AETX, △: 1 µM AETX, ▽: 2 µM AETX, ◇: 5 µM AETX, ◁: 10 µM AETX, ▷: 30 µM AETX. Addition of 1: AETX, 2: oligomycin, 3: FCCP, 4: rotenone/antimycin A + Hoechst 33342. Whiskers indicate standard deviation (1x).

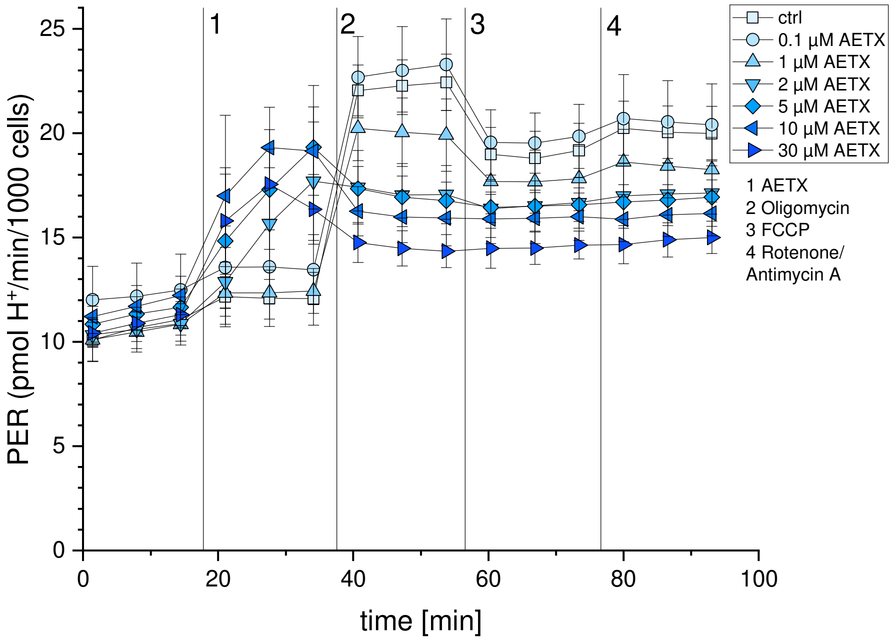

**Fig. S16:** Proton Efflux Rate (PER) measurement of HeLa cells after acute stimulation with AETX. □: solvent control (0.1% DMSO), ○: 0.1 µM AETX, △: 1 µM AETX, ▽: 2 µM AETX, ◇: 5 µM AETX, ◁: 10 µM AETX, ▷: 30 µM AETX. Addition of 1: AETX, 2: oligomycin, 3: FCCP, 4: rotenone/antimycin A + Hoechst 33342. Whiskers indicate standard deviation (1x).

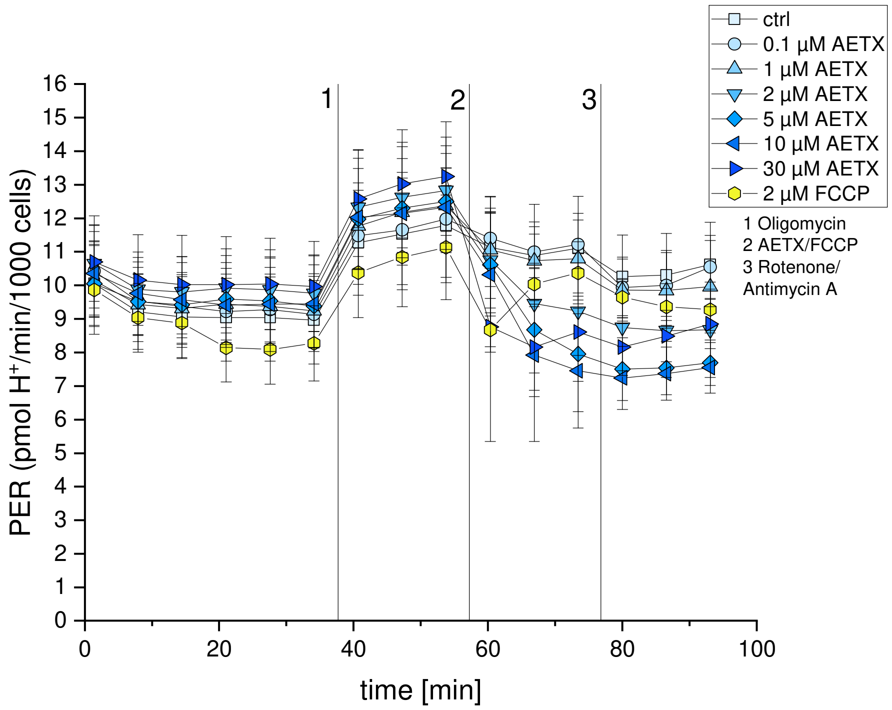

**Fig. S17:** Proton Efflux Rate (PER) measurement of fibroblasts after acute stimulation with AETX or FCCP subsequent to FoF1-ATPase inhibition. □: solvent control (0.1% DMSO), ○: 0.1 µM AETX, △: 1 µM AETX, ▽: 2 µM AETX, ◇: 5 µM AETX, ◁: 10 µM AETX, ▷: 30 µM AETX, ⬡: 2 µM FCCP. Addition of 1: oligomycin, 2: AETX or 2 µM FCCP, 3: rotenone/antimycin A + Hoechst 33342. Whiskers indicate standard deviation (1x).

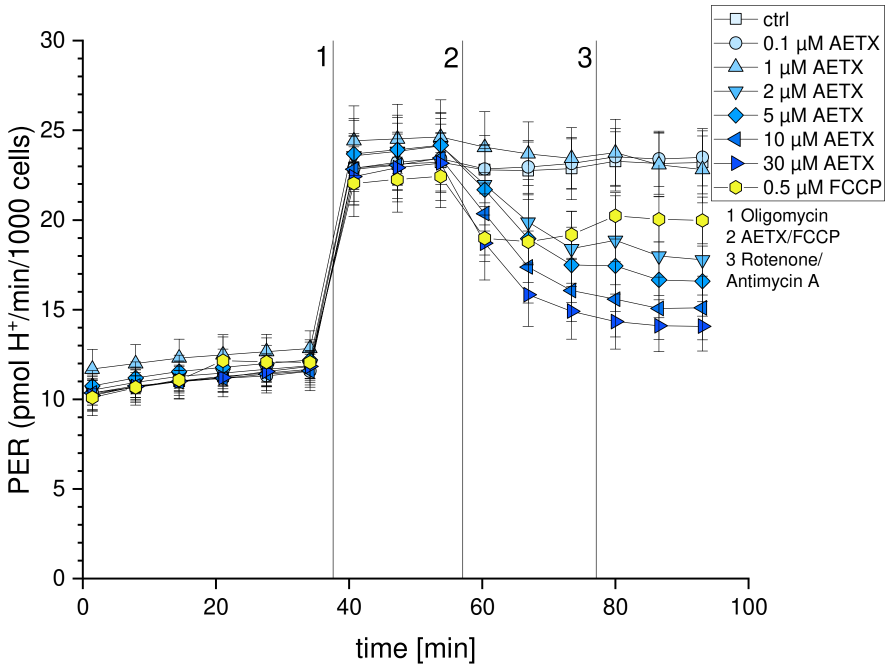

**Fig. S18:** Proton Efflux Rate (PER) measurement of HeLa cells after acute stimulation with AETX or FCCP subsequent to FoF1-ATPase inhibition. □: solvent control (0.1% DMSO), ○: 0.1 µM AETX, △: 1 µM AETX, ▽: 2 µM AETX, ◇: 5 µM AETX, ◁: 10 µM AETX, ▷: 30 µM AETX, ⬡: 0.5 µM FCCP. Addition of 1: oligomycin, 2: AETX or 0.5 µM FCCP, 3: rotenone/antimycin A + Hoechst 33342. Whiskers indicate standard deviation (1x).

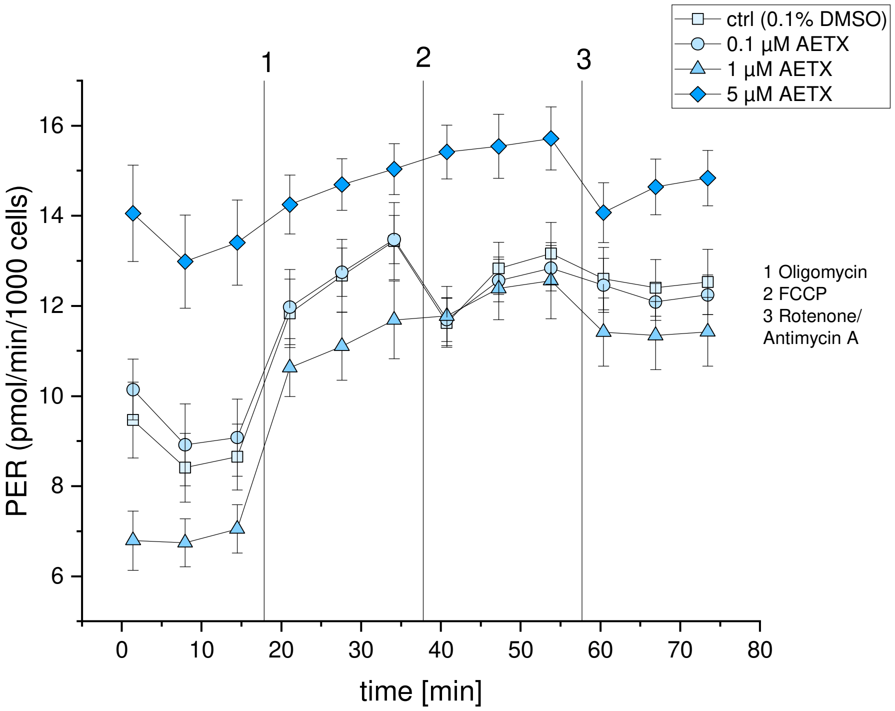

**Fig. S19:** Proton Efflux Rate (PER) measurement of fibroblasts after 24 h of treatment with AETX. □: solvent control (0.1% DMSO), ○: 0.1 µM AETX, △: 1 µM AETX, ◇: 5 µM AETX. Addition of 1: oligomycin, 2: FCCP, 3: rotenone/antimycin A + Hoechst 33342. Whiskers indicate standard deviation (1x).

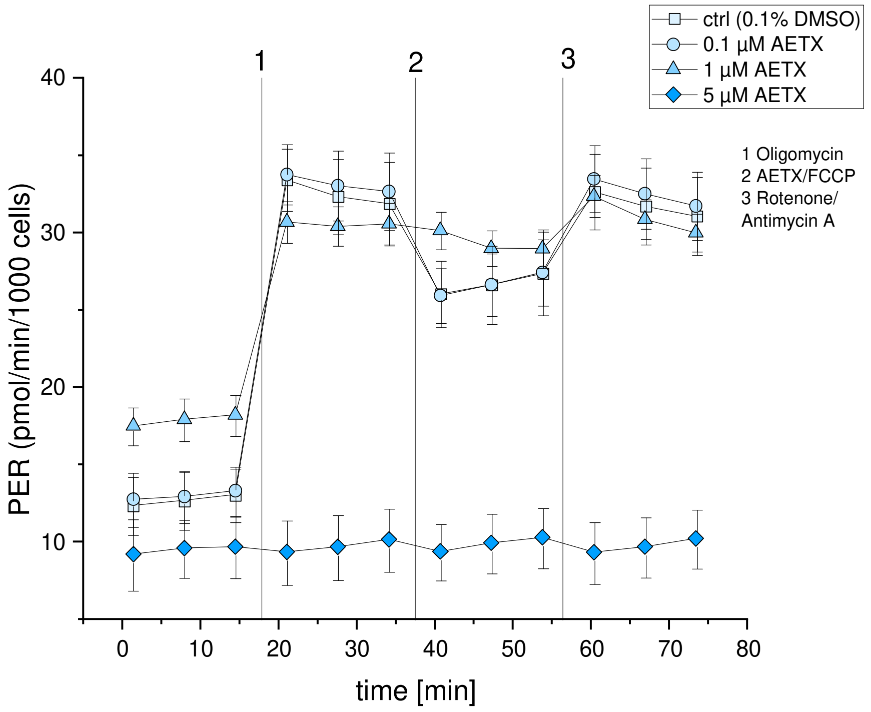

**Fig. S20:** Proton Efflux Rate (PER) measurement of HeLa cells after 24 h of treatment with AETX. □: solvent control (0.1% DMSO), ○: 0.1 µM AETX, △: 1 µM AETX, ◇: 5 µM AETX. Addition of 1: oligomycin, 2: FCCP, 3: rotenone/antimycin A + Hoechst 33342. Whiskers indicate standard deviation (1x).

**Proton leak and coupling efficiency**

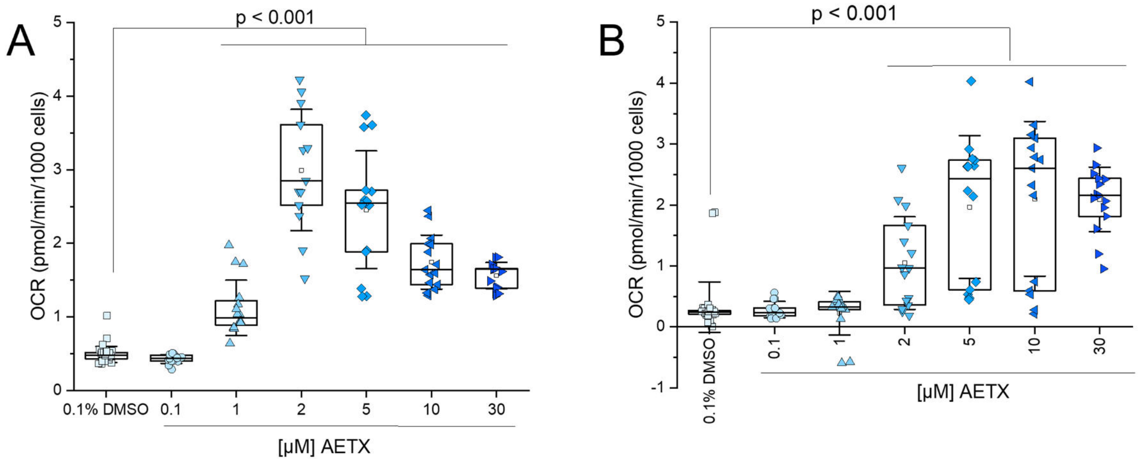

**Fig. S21:** Proton leak in **(A)** HeLa cells, and **(B)** fibroblasts, both treated with AETX. □: solvent control (0.1% DMSO), ○: 0.1 µM AETX, △: 1 µM AETX, ▽: 2 µM AETX, ◇: 5 µM AETX, ◁: 10 µM AETX, ▷: 30 µM AETX. Statistically significant difference to control obtained with Student‘s t-test. Whiskers indicate standard deviation (1x). Boxes indicate 25-75 percentile with the median as line and mean as open square.

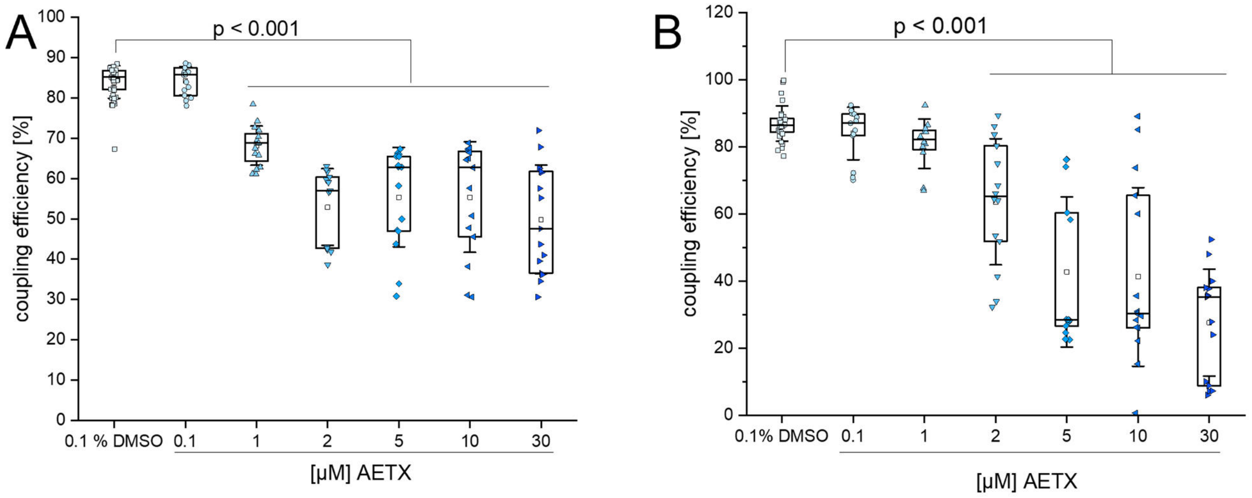

**Fig. S22:** Coupling efficiency in **(A)** HeLa cells, and **(B)** fibroblasts, both treated with AETX. □: solvent control (0.1% DMSO), ○: 0.1 µM AETX, △: 1 µM AETX, ▽: 2 µM AETX, ◇: 5 µM AETX, ◁: 10 µM AETX, ▷: 30 µM AETX. Statistically significant difference to control obtained with Student‘s t-test. Whiskers indicate standard deviation (1x). Boxes indicate 25-75 percentile with the median as line and mean as open square.

**Supporting OCR**

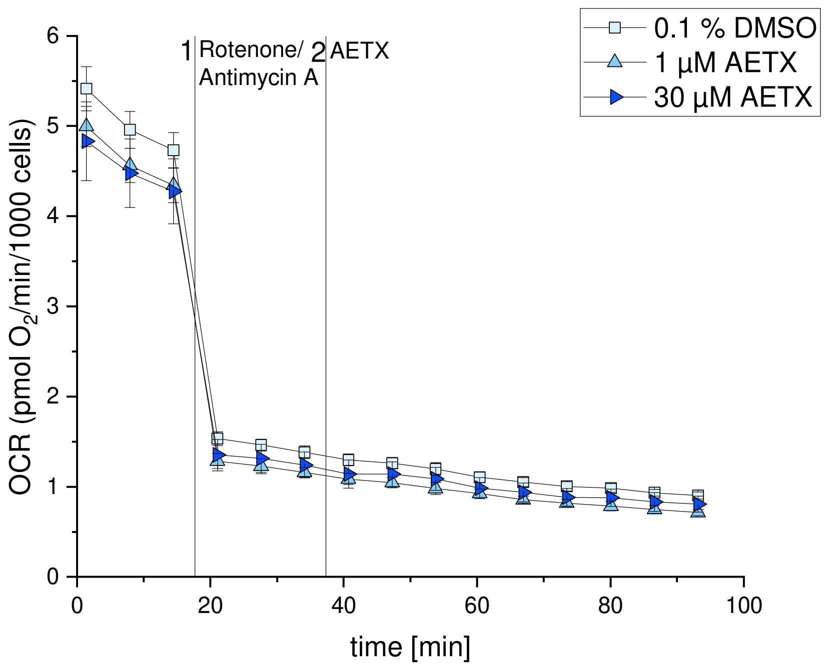

**Fig. S23:** Oxygen consumption rate (OCR) in HeLa cells after inhibition of the respiratory chain with rotenone/ Antimycin A (1). After the inhibition, AETX was added (2). □: solvent control (0.1% DMSO), △: 1 µM AETX, ▷: 30 µM AETX. Cave: biological replicate = 1.

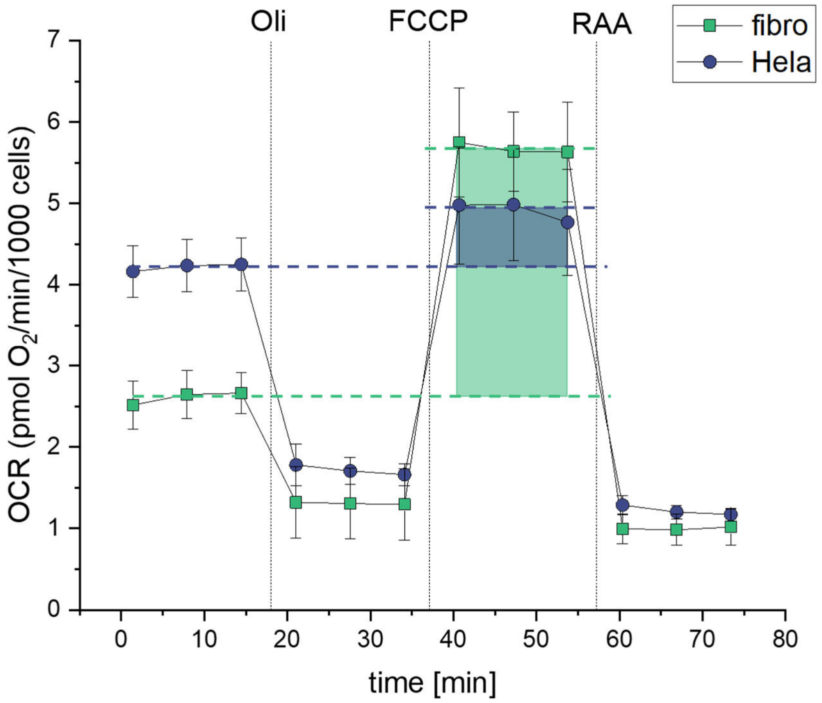

**Fig. S24:** Mean oxygen consumption rate (OCR) of fibroblasts (fibro, green □) and HeLa cells (Hela, blue ○). Cells were treated with 0.1% DMSO and subsequently with oligomycin (Oli), FCCP and rotenone/antimycin A + Hoechst 33342 (RAA). Whiskers indicate standard deviation (1x). Spare respiratory capacity indicated as colored areas, green: fibroblasts, blue: HeLa cells.

**Cell biology – enhanced cytotoxicity**

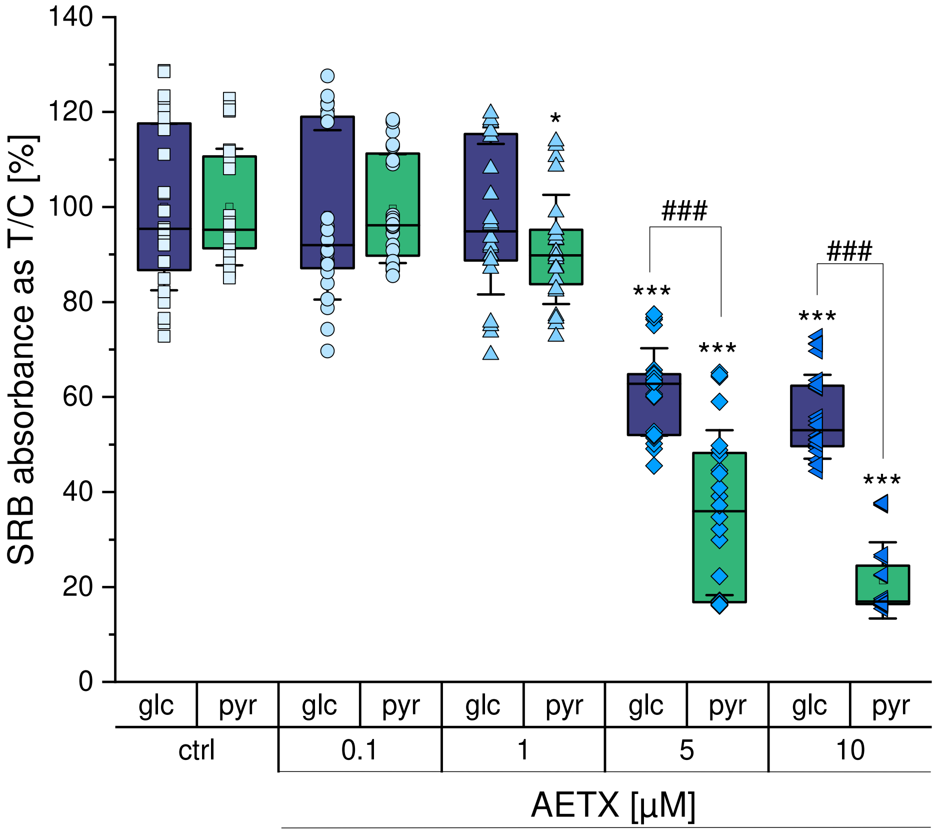

**Fig. S25:** Mean SRB absorbance of AETX-treated HCT116 cells in dependence of different media supply. Data is normalized to the control. Green: pyruvate containing medium (pyr), violet: glucose containing medium (glc). □: 0.1% DMSO (solvent control), ○: 0.1 µM, △: 1 µM, ◇: 5 µM, ◁: 10 µM. Boxes indicate 25-75 percentile with the median as line. Standard deviation (1x) indicated as whiskers. ### p < 0.001 comparison between the different media, */ *** p < 0.05/0.001 comparison to the respective solvent control. Statistical results obtained with Mann-Whitney test. Data from six independent biological replicates.

### Cell biology – ROS

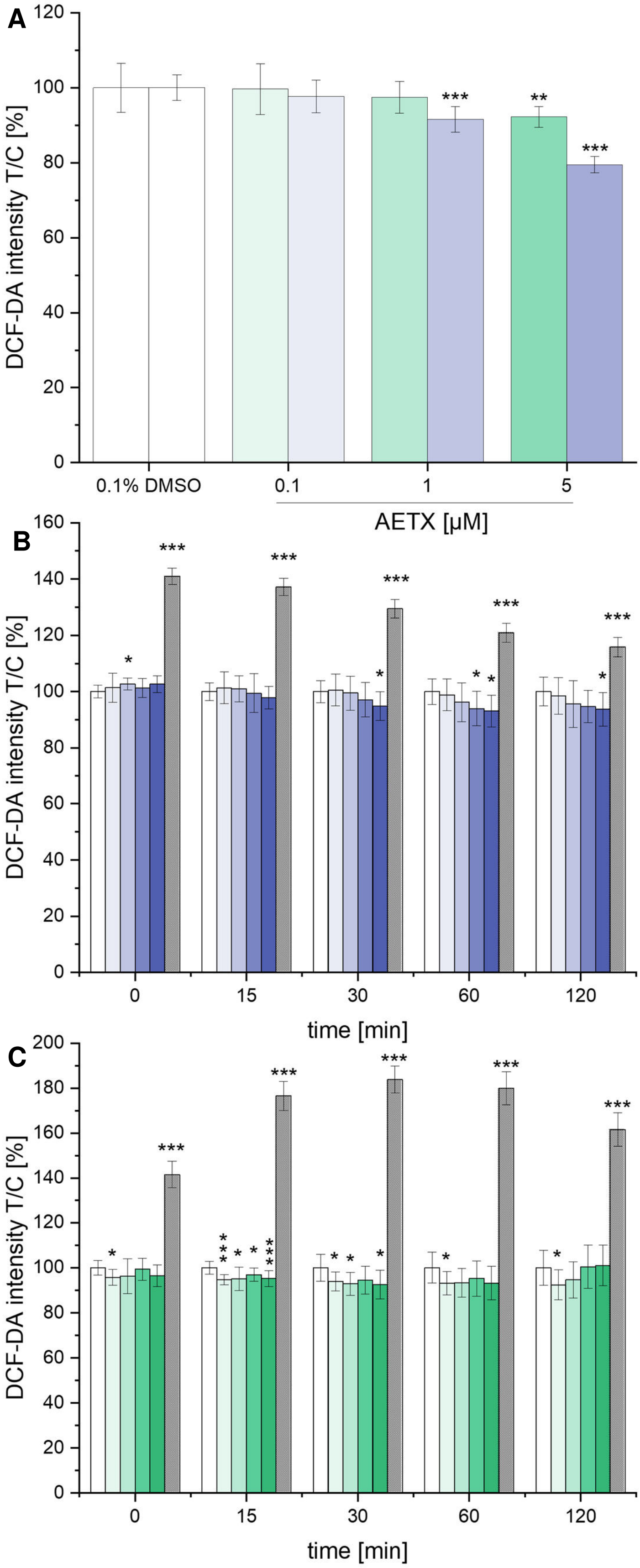

**Fig. S26:** Effect of AETX on ROS formation detected with DCF-DA. **(A)** Mean DCF-DA intensity of AETX-treated Hela cells and fibroblasts after 24 h incubation normalized to control. **(B)** Kinetic of mean DCF-DA intensity of AETX-treated HeLa cells, and **(C)** fibroblasts normalized to the control. Grey bars: positive control ($H_2O_2$). Panel **A – C**: violet: HeLa cells, green: fibroblasts. Increasing shades indicate increasing concentrations of AETX (0.1% DMSO (control), 0.1, 1, 5, and 10 µM AETX); Statistically significant difference to the control indicated with */**/*** $p < 0.05/ 0.01/ 0.001$, obtained with Student's t-test. Whiskers indicate standard deviation (1x).

### Chemistry – purity of AETX, dn-AETX, m-AETX

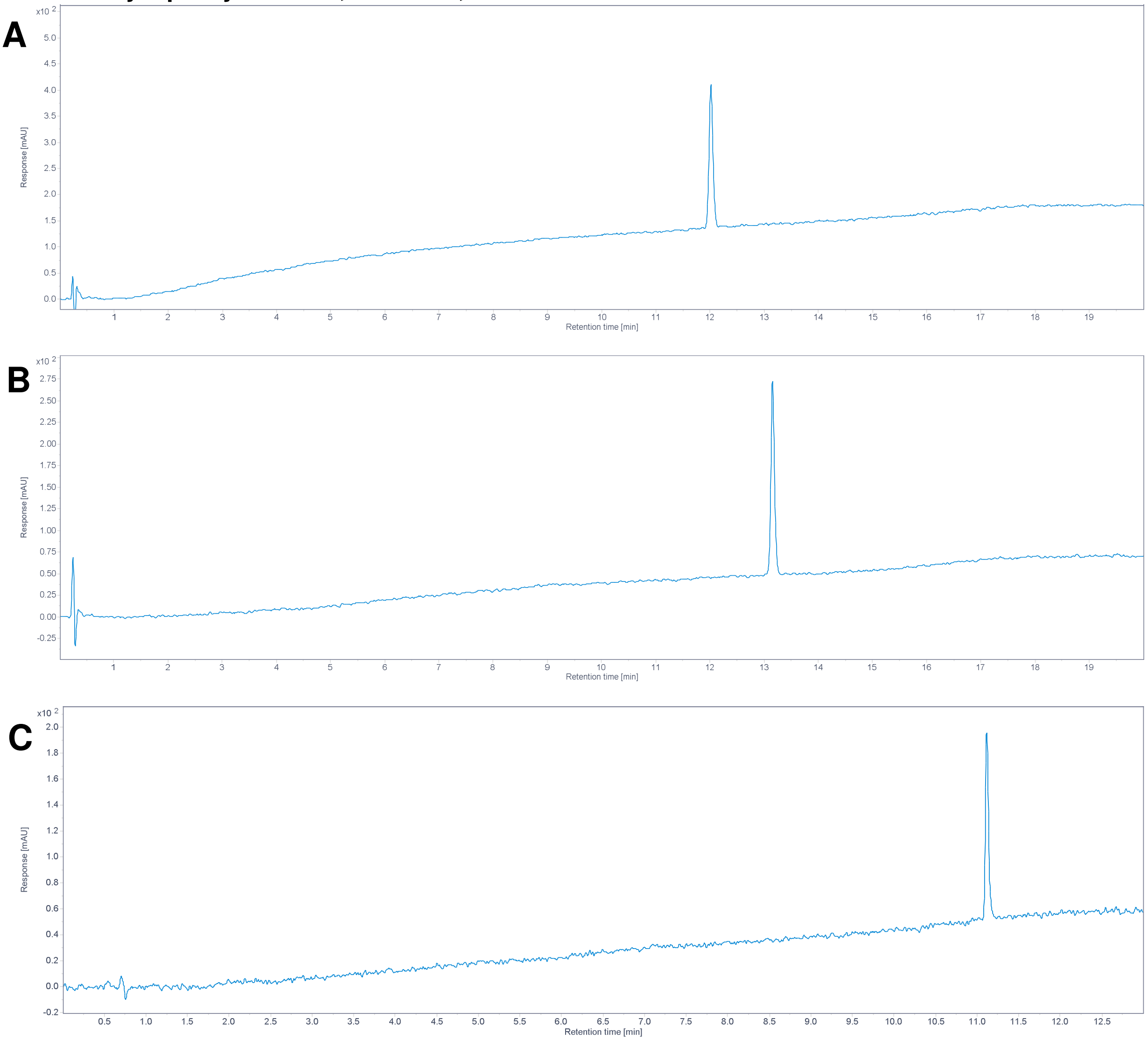

**Fig. S27:** HPLC-UV chromatogram of AETX **(A)**, dn-AETX **(B)** and m-AETX **(C)** at 210 nm. Purity of all compounds >99.5%.

### Chemistry – structure confirmation of m-AETX

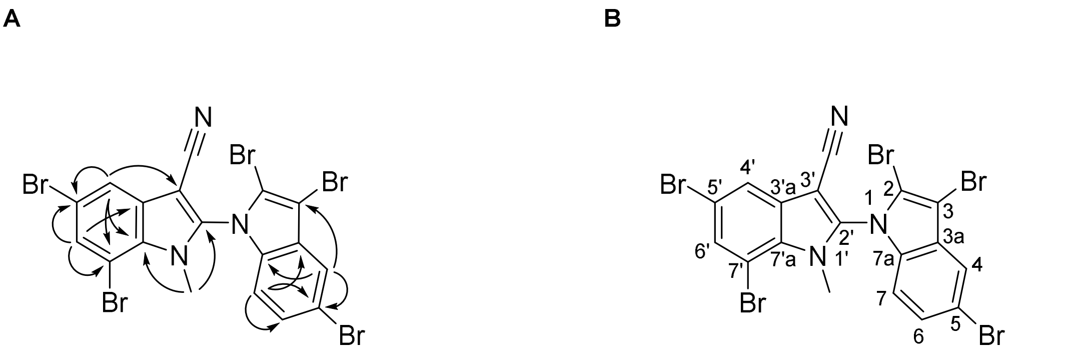

2,3,5,5',7'-pentabromo-1'-methyl-1'*H*-[1,2'-biindole]-3'-carbonitrile

**Fig. S28:** Structure confirmation with **(A)** key HMBC correlations, **(B)** numbering and IUPAC name.

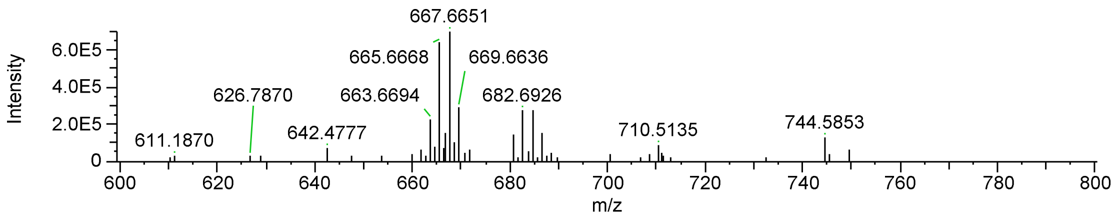

**Fig. S29:** HRMS spectrum of m-AETX (pos. mode). $[M+H]^+$ at *m/z* 667.6651, $C_{18}H_9N_3{}^{79}Br_2{}^{81}Br_3$ (calc.667.6652, Δ 0.1 ppm)

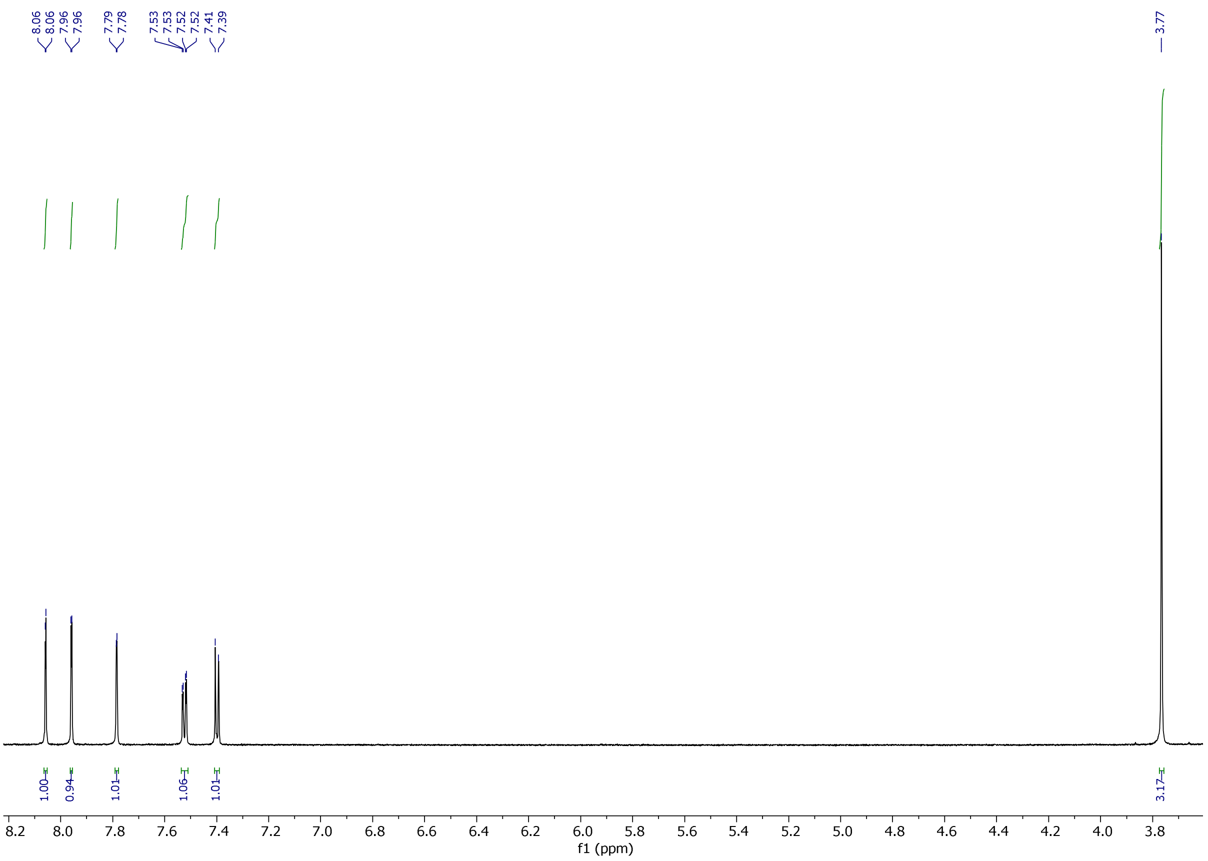

**Fig. S30:** $^1H$ NMR spectrum (700 MHz) of m-AETX in DMSO-$d_6$ (3.5 to 8.2 ppm).

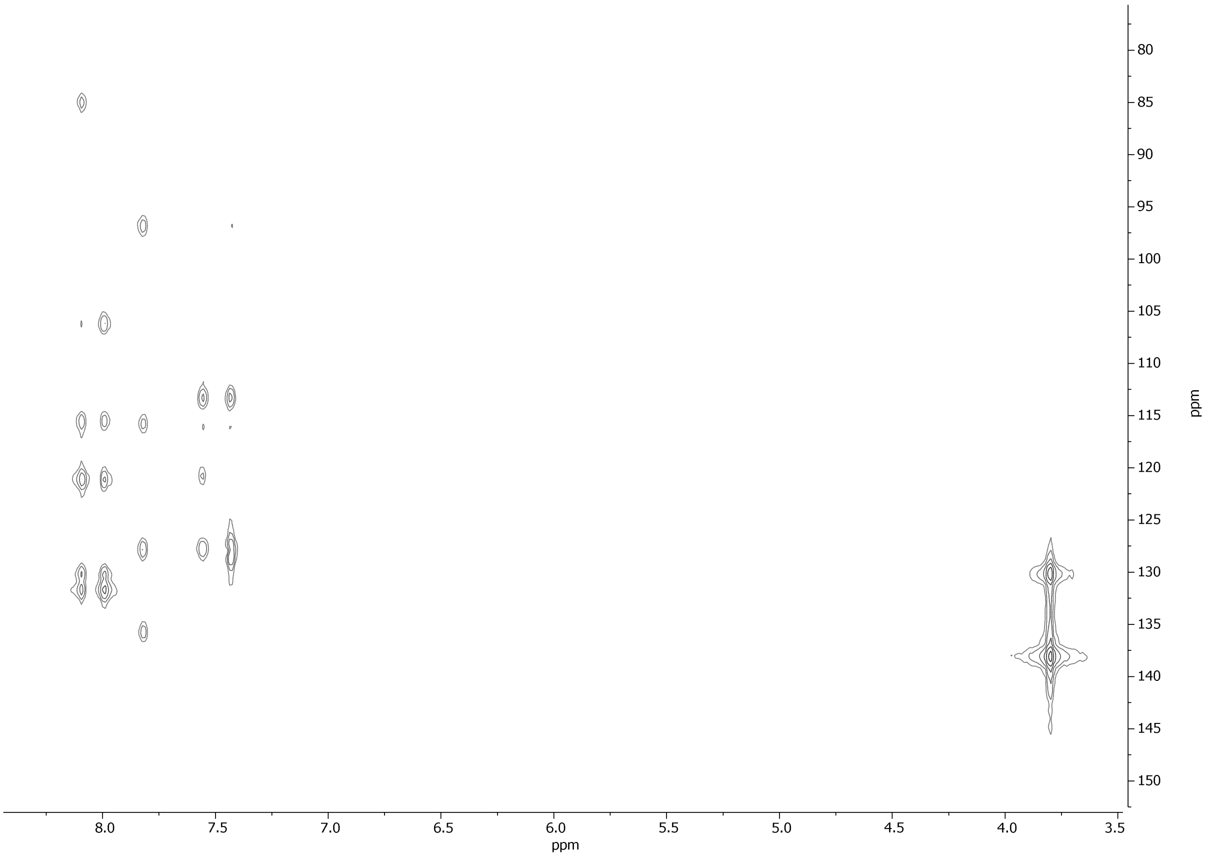

**Fig. S31:** $^{13}C$-HMBC NMR spectrum (700 MHz) of m-AETX in DMSO-$d_6$ (3.4 to 8.4 ppm).

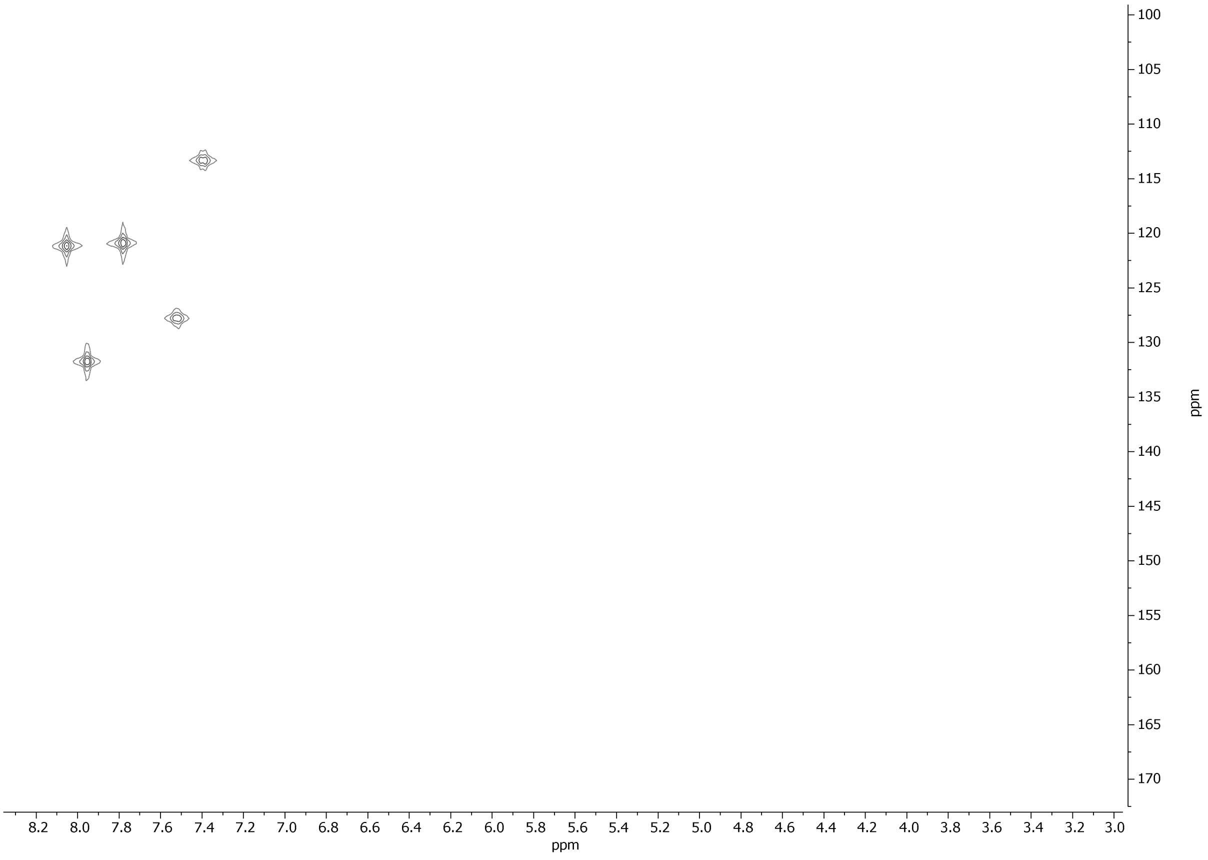

**Fig. S32:** $^{13}C$-HSQC NMR spectrum (700 MHz) of m-AETX in DMSO-$d_6$ (3.0 to 8.2 ppm).
